# Age-dependent brain pigmentation drives early neuroinflammatory molecular signatures linked to neurodegeneration

**DOI:** 10.64898/2026.08.03.742448

**Authors:** Núria Peñuelas, Helena Xicoy, Marina Lorente-Picón, Alba Nicolau-Vera, Annabelle Parent, Marta Gonzalez-Sepulveda, Javier Hoyo, Ariadna Laguna, Miquel Vila

## Abstract

**Background:** Neuromelanin (NM) is a pigment that progressively accumulates with age in catecholaminergic neurons, particularly in the substantia nigra, ventral tegmental area, and locus coeruleus. These neuronal populations are especially vulnerable to degeneration in Parkinson’s disease (PD). Elevated intracellular NM levels have been linked to neurodegeneration and PD-like phenotypes in experimental models. However, the molecular mechanisms underlying NM-induced pathology remain poorly understood, as human studies cannot disentangle the specific effects of NM accumulation from those of normal aging.

**Methods:** We performed transcriptomic microarray analysis on laser-captured catecholaminergic neurons and regions (substantia nigra, ventral tegmental area, locus coeruleus) from NM-producing transgenic mice (tgNM) and NM-free wild-type controls across different ages, and compared them to data from postmortem human brain tissue. One of the molecular targets identified, GPNMB, was validated in mouse and human tissue, and functionally tested *in vivo*.

**Results:** We identified region- and age-dependent transcriptional changes associated with progressive NM accumulation. NM consistently upregulated neuroinflammatory pathways with enrichment of disease-associated microglial genes, while downregulating transcription, translation, and mitochondrial functions. Locus coeruleus exhibited the earliest and strongest transcriptional alterations, whereas substantia nigra and ventral tegmental area showed a later-onset, age-progressive transcriptional dysfunction. Neuron-specific analyses revealed that many changes originated within NM-containing neurons rather than being solely glial-driven. NM-driven transcriptional profiles in mice strongly correlated with postmortem data from PD patients, underscoring their translational relevance. Among molecular targets, the glycoprotein GPNMB was consistently upregulated in NM-containing neurons and validated at RNA and protein levels in both NM-producing transgenic mice and in human PD brains. Functional experiments demonstrated that GPNMB overexpression attenuated NM-linked dopaminergic neurodegeneration and improved motor performance in mice.

**Conclusion:** This study provides a comprehensive *in vivo* characterization of NM-specific transcriptomic changes in catecholaminergic neurons, showing that NM accumulation drives neuroinflammatory and neurodegenerative programs. Our results support that the neuroinflammatory changes observed in tgNM mice and in human PD represent early pathological events that precede overt neurodegeneration. The disease-associated gene GPNMB emerged as a conserved NM-induced factor with protective properties, highlighting its potential as a therapeutic target in PD and aging-related neurodegeneration.

## Introduction

Neuromelanin (NM) pigment in the human brain appears at the age of three years old and gradually accumulates with age in all catecholaminergic neuronal groups A1-A14.^1–4^ The most heavily pigmented catecholaminergic neurons in the human brain constitute the most vulnerable areas in Parkinson’s disease (PD): the dopaminergic neurons in the substantia nigra pas compacta (SN) and the ventral tegmental area (VTA), and the noradrenergic neurons in the locus coeruleus (LC).^5–7^ This observation points to NM as a potential key factor involved in the specific regional vulnerability occurring in PD. However, this hypothesis remained speculative for a long time as NM accumulation in the human brain coincides with aging processes that affect a broad range of cellular functions, obscuring the exact role of NM in physiological aging. Additionally, this hypothesis could not be experimentally addressed since most animal species used as *in vivo* models lack this pigment.^5,8^

NM is now emerging as a new player in the etiology of PD due to newly developed experimental animal models that reproduce human-like NM accumulation in neuronal populations^9^. These NM-producing experimental *in vivo* models overexpress the human melanogenic enzyme tyrosinase (TYR) either through adenoassociated viral vectors (AAV) into a specific brain area^10,11^ or through classical transgenesis in constitutive tissue-specific TYR-expressing mice^12^. Specifically, TYR overexpression through AAV in the mouse SN leads to intracellular NM accumulation at levels comparable to those observed in postmortem PD SN neurons, resulting in Lewy body (LB)-like inclusion body formation, neuroinflammation, nigrostriatal neurodegeneration, and motor deficits^11^. Conversely, TYR overexpression in the mouse LC induces noradrenergic neurodegeneration linked to changes in sleep patterns and anxiety-like behavior^10^. Recently, our group has developed and characterized a new transgenic mouse model (referred to as tgNM) that overexpresses TYR under the tyrosine hydroxylase (TH) promoter, reproducing age-dependent neuronal pigmentation in a brain-wide manner (i.e. following the distribution of catecholaminergic groups in the human brain)^12^. This new NM-producing model shows features of prodromal PD, including dopaminergic dysfunction, noradrenergic neurodegeneration, LB-like inclusion body formation, and neuroinflammatory changes in all pigmented areas in the brain, which are associated with both motor and non-motor PD-like phenotypes^12^.

The precise molecular consequences of age-dependent NM accumulation have largely remained elusive due to the absence of suitable model systems. Numerous studies have characterized the SN in postmortem brain tissues of PD patients compared to controls using bulk dissections^13–20^ or single-cell preparations.^21–24^ Conversely, there are few studies characterizing the LC in PD compared to controls.^25,26^ While these studies define the PD postmortem transcriptomic signature, they do not address the potential contribution of age-dependent NM accumulation *per se* to the PD neurodegenerative phenotype, as healthy control human brains (i.e. non-PD) also gradually accumulate NM with aging. To overcome these limitations, here we first characterized the tgNM mouse model across brain regions and ages to define NM-associated transcriptional changes in vivo, comparing age-matched neuronal phenotypic groups in the presence or absence of NM, and assessing the intrinsic susceptibility of different catecholaminergic cell types to progressive, age-dependent NM accumulation. In addition, we used AAV-mediated TYR overexpression as an independent experimental approach to functionally assess the impact of the NM-induced gene GPNMB on dopaminergic neurodegeneration in a TYR-driven NM-producing model.

## Methods

This study complies with all relevant ethical regulations. All procedures involving human samples were conducted in accordance with guidelines established by the BPC (CPMP/ICH/135/95) and the Spanish regulation (223/2004) and approved by the Vall d’Hebron Research Institute (VHIR) Ethical Clinical Investigation Committee (PR(AG)370/2014). All the experimental and surgical procedures using animals were conducted in strict accordance with the European (Directive 2010/63/UE) and Spanish laws and regulations (Real Decreto 53/2013; Generalitat de Catalunya Decret 214/97) on the protection of animals used for experimental and other scientific purposes. Protocols were approved by the Vall d’Hebron Research Institute (VHIR) Ethical Experimentation Committee and the Generalitat de Catalunya (Protocol 11442).

### Human postmortem brain tissue

Fresh frozen and paraffin-embedded human postmortem brain tissue were obtained to evaluate GPNMB expression. Samples from neurologically healthy control individuals (n=13) and from patients diagnosed with idiopathic Parkinson’s disease (iPD) or Parkinson’s disease dementia (PDD) (n=14) were provided by the Neurological Tissue BioBanks at IDIBAPS–Hospital Clínic and Vall d’Hebron Hospital (Barcelona) and the Biobanco en Red de la Región de Murcia (BIOBANC-MUR). The mean age (±SD) was 80.00 ± 7.9 years for controls and 78.50 ± 6.2 years for iPD/PDD cases. The average postmortem interval (PMI) was 8.18 ± 4.2 hours. A detailed description of all human samples used in the study is provided in the submitted Supplementary Data 5.

### TgNM mouse colony

Transgenic mice B6.Cg-Tg(Th-TYR)26Mvila/Mvila (RRID:MGI:7730751) were obtained by pronuclear microinjection of the human tyrosinase complementary DNA (cDNA) fused to the rat tyrosine hydroxylase promoter into C57Bl6-SJL mouse zygotes as described elsewhere ^12^. Mice were backcrossed for 8– 10 generations using C57BL/6 J mice (Charles River; RRID:MGI:3028467) and maintained as heterozygotes. Animals were housed two to five per cage with ad libitum access to food and water during a 12-hour (h) light/dark cycle (light-on 8 a.m.). Mice were randomly distributed into the different experimental groups and control and experimental groups were processed at once to minimize bias. Both male and female mice were included in the study. In transcriptomic and histological experiments, sexes were evenly distributed across experimental groups to avoid sex bias. Sex was not treated as a primary biological variable in the experimental design due to power constraints and the need to reduce the total number of animals used. Accordingly, no sex-specific subgroup analyses were performed.

### Brain processing for LCM

Mice (tgNM and wt) aged 3, 12 and 20 months (m) (n=6 animals per genotype and age) were sacrificed by cervical dislocation. Brains were removed, snap-frozen for 20s in dry-iced cooled 2-methylbutane (isopentane) and stored at −80°C until further processing. Before sectioning, mouse and human brain samples were tempered from −80°C to −20°C for 1h and then sectioned in a cryostat (CM3050S Leica, Germany). 10µm sections from postmortem SN tissue or serial 10µm sections covering the whole rostro-caudal extent of the mouse SN and VTA regions (every 12th section for a total of 6-7 sections/animal) were collected on specialized membrane-coated slides for LCM (PEN-membrane 2,0um, MDG3P40W, MicroDissect GmbH), previously irradiated with UV light for 1h to reduce static electricity. Slides were kept at −20°C during sectioning and then stored at −80°C until processing.

### Histogene staining for LCM

Sections were stained following the Arcturus® HistoGene® Frozen Section Staining Kit (#0401, Thermo Fisher Scientific). Briefly, slides were transferred on dry ice from −80°C immediately to a series of slide staining jars containing RNase-free solutions. Slides were kept in 75% ethanol for 30s, H_2_0 for 30s, HistoGene® solution (100 µl) for 1min and finally dehydrated with increasing concentrations of RNase-free ethanol solutions (H_2_0-15s, ethanol 75%-15s; 95%-15s; 100%-15s). In order to reuse staining jars, a rigorous cleaning procedure was performed between experiments. When reused, jars were washed twice with 100% ethanol, washed twice with distilled H_2_0, kept in Molecular BioProducts RNase away surface decontaminant solution (ThermoScientific, #7000TS1) overnight, washed three times with autoclaved miliQ H_2_0 for and dried under a fume hood for a complete removal of remaining solutions.

### Laser capture microdissection (LCM)

To ensure accurate identification and rapid collection of neuronal groups, catecholaminergic regions were delineated for each animal/human specimen in rostrocaudal 10µm serial sections immunostained with TH following standard immunohistochemical protocols after quenching cryosections for 10 min in 3% H2O2-10% (vol/vol) methanol (see GPNMB validation/Immunohistochemistry). Subsequently, neurons/regions were rapidly collected from consecutive serial 10µm sections stained with the nonspecific, fast-penetrating stain Histogene (Fig. 1). Immediately after the histogene staining procedure, the slides were placed under the laser capture microdissection microscope (Leica LMD6500), previously decontaminated with Molecular BioProducts RNase away surface decontaminant (ThermoScientific, #7000TS1). Neurons/regions of interest were visualized with the Leica LMD software (Leica Laser Microdissection V6.6.2), drawn manually using a 40x/5x objective, cut with a UV-laser and collected into an autoclaved RNase/DNase Free eppendorf cap (0.65 ml Low Binding MCT, Sorenson Biosciences, #11300) containing 30µl of QIAazol Lysis Reagent solution (from miRNeasy Micro Kit, Qiagen, #217084). Approximately 300 neurons/10 regions were collected from each region and animal/individual. The total time of the LCM procedure never exceeded 1h per slide to avoid compromising the quality of the tissue RNA. Eppendorf tubes containing collected samples were vortexed to ensure complete lysis and immediately frozen on dry ice. Samples were stored at −80°C for subsequent RNA isolation.

**Figure 1.**
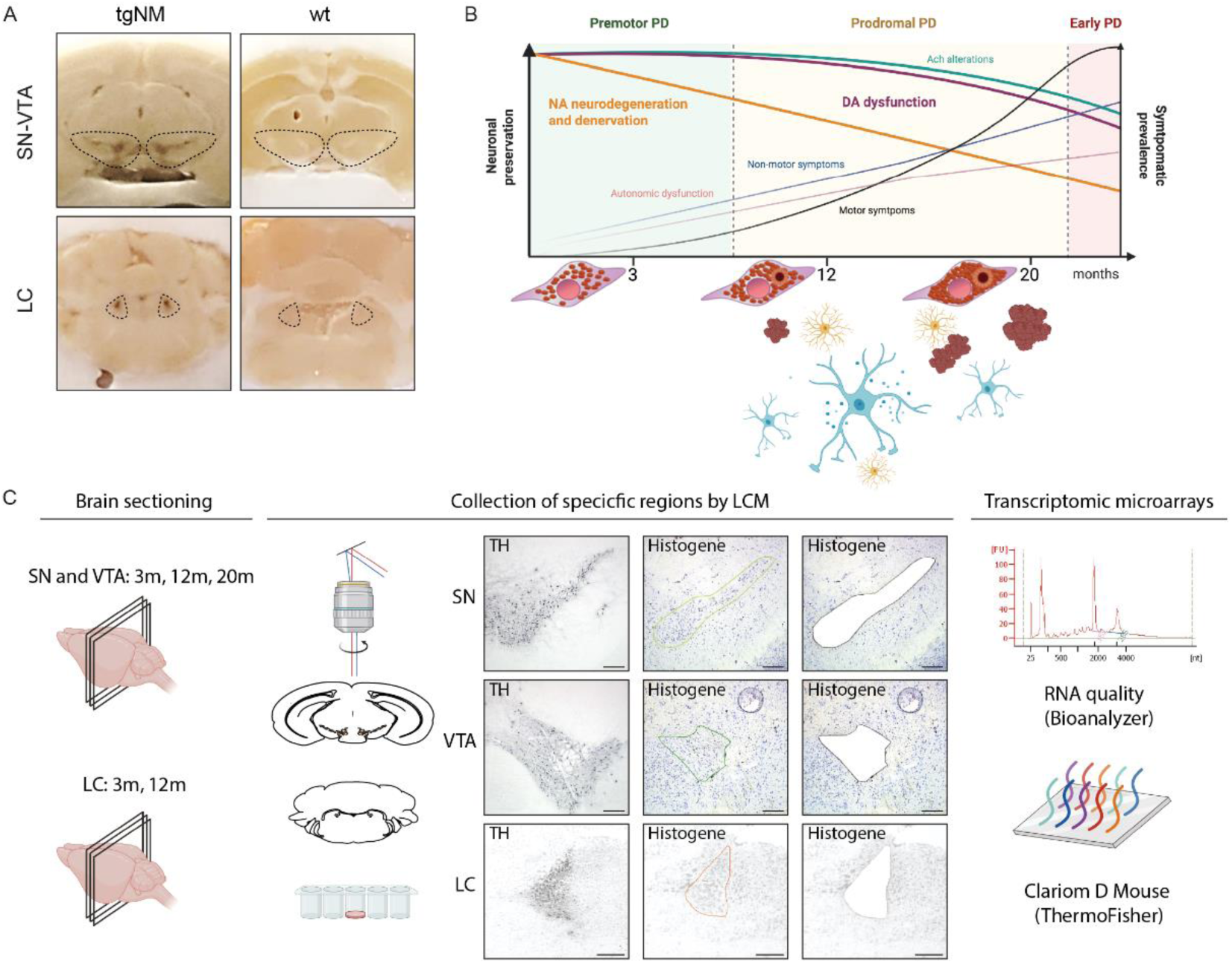
Age-dependent neurodegeneration and transcriptomic profiling of catecholaminergic neuromelanin-pigmented brain regions in tgNM mice. (a) Representative coronal brain sections showing neuromelanin (NM) accumulation in tgNM mice compared to wild-type (wt) littermates in the substantia nigra (SN) and ventral tegmental area (VTA) (upper panels), and in the locus coeruleus (LC) (lower panels). Dotted lines delineate the anatomically defined regions of interest. Adapted from Laguna, Peñuelas et al. ^12^. (b) Schematic illustration of the progressive pathological changes observed in tgNM mice, highlighting the temporal dynamics of noradrenergic (NA) neurodegeneration, dopaminergic (DA) and cholinergic (Ach) dysfunction, and of motor and non-motor behavioral phenotypes associated with Parkinson’s disease (PD). The model also recapitulates hallmark PD features including α-synuclein-positive Lewy body-like inclusions, extraneuronal NM granules, and neuroinflammatory responses. Key time points (3, 12, and 20 months) correspond to increasing NM accumulation and increased regional vulnerability in tgNM mice. (c) Workflow of the experimental design for transcriptomic profiling. Left: Brain sectioning strategy showing the anatomical levels sampled for SN and VTA (3, 12, and 20 months) and LC (3 and 12 months). Middle: Catecholaminergic nuclei were identified by tyrosine hydroxylase (TH) immunostaining, and target regions were selectively isolated using laser capture microdissection (LCM) on adjacent Histogene-stained 10 µm sections. Representative images illustrate regions before and after LCM. Right: Total RNA was extracted from the microdissected tissue, quality assessed via Bioanalyzer electropherogram, and subsequently analyzed using Clariom D Mouse transcriptomic microarrays (ThermoFisher). Scale bar: 200µm. months, m.

### RNA extraction and quality control

Isolation of total RNA from the microdissected tissue and neurons was performed using the miRNeasy Micro Kit (Qiagen, #217084). This procedure combines phenol/guanidine-based lysis of samples and silica-membrane– based purification of total RNA, including long and small RNA species. Samples kept at −80°C after the LCM procedure were quickly thawed and QIAazol Lysis Reagent was added up to a final volume of 152 µl. Samples were vortexed for further lysis of tissue and cells, left for 5min at room temperature to promote dissociation of nucleoproteins, and then, chloroform was added to samples and incubated for 2-3min at room temperature and centrifuged to allow phase separation. Next, the aqueous phase was transferred to a new tube, mixed with 100% ethanol and passed through an RNeasy MinElute spin column. The column membrane was washed twice with wash buffers provided by the kit and with 80% ethanol, and finally, RNA was eluted with RNase-free water (14 µl) and stored at −80°C for its subsequent analysis.

RNA concentration and quality were assessed by electrophoresis using an Agilent 2100 BioAnalyzer instrument (Agilent technologies) and an Agilent RNA 6000 Pico Kit (0,05-5ng/ul) (Agilent technologies). Only samples with an RNA integrity number (RIN)>5 (or RIN>4 for human LCM samples) on a scale from 1 (highly degraded) to 10 (highest integrity) were processed for subsequent analysis (RIN mean±SD; tgNM SN region: 6.43±0.8, tgNM VTA region: 7.01±1.1, tgNM LC region:7.4±0.2, tgNM SN neurons: 6.5±1, tgNM VTA neurons: 6.4±0.9, human SN region=6.01±1.2, human LC region: 6.32±0.9, human FC region=6.79±1.0, human SN neurons: 4.64±1.1). RNA quality did not differ between experimental groups nor correlate with time spent during the LCM collection process, confirming that the protocol for isolating pigmented areas was not compromising RNA quality.

### Transcriptomic microarrays

RNA samples extracted from tgNM and wt animals aged 3m, 12m and 20m that passed the RNA quality control (n=4-6 mice per genotype/region/age) were analyzed using whole-transcriptome microarrays Clariom D Pico Assay, mouse (#902664) and washing kit GenChip Hyb Wash & Stain kit (#900720) (Applied Biosystems, ThermoFischer Scientific) [n=32 (SN neurons), n=30 (VTA neurons), n=24 (LC region), n=30 (SN region) and n=36 (VTA region)] in the High Technology Unit from Vall d’Hebron Research Institute (VHIR-UAT). Each region was processed in independent microarray experiments and both genotype and age variables were distributed evenly in the different experimental batches. Clariom D mouse microarrays assess >214,000 transcripts mapping to both coding and non-coding transcripts. RNA reverse transcription is initiated at the poly-A tail as well as throughout the entire length of the RNA molecule enabling the amplification of intact, partially degraded, and compromised RNA samples (starting from 100pg of RNA).

### Differential expression analysis (DEA) and Biological Significance Analysis (BSA)

Normalization and analysis of the raw data was performed by the Unit of Statistics and Bioinformatics (UEB) from the Vall d’Hebron Research Institute (VHIR). Arrays were considered outliers when they consistently deviated from the main cohort in at least two independent quality control analyses (inspection of raw intensity distributions, Principal Component Analysis (PCA), hierarchical clustering, and heatmap visualization), and were consequently removed from downstream analyses [n=3 SN neurons, n=3 VTA neurons, n=1 SN region]. During the pre-processing and normalization step, a slight batch effect was detected in all experimental groups and thus incorporated in the differential expression analysis. Sex effect was also detected in some experimental groups and thus, also incorporated in the analysis (SN, VTA, LC regions). Differential expression analysis (DEA) between pigmented and non-pigmented areas was performed using R software and the libraries developed for the microarray analysis in the Bioconductor Project ^27^. DEA was based on adjusting a linear model with empirical Bayes moderation of the variance, a technique similar to ANOVA specifically developed for microarray data analysis by Gordon K Smyth^28^. To identify statistically significant differentially expressed genes (DEGs) a threshold of raw p-value <0.01 and FC>|1.5| was applied (Supplementary Data 1). Biological Significance Analysis (BSA) was performed using the gene set enrichment analysis (GSEA) function and the *Gene-Ontology (GO)* and *Reactome* databases (Supplementary Data 2). A correction for multiple comparisons was applied for statistical significance [False Discovery Rate (FDR) threshold <0.05]. To reduce redundancy and enhance interpretability of pathway enrichment results, we summarized enriched terms using the simplifyEnrichment R package v1.6.1^29^ (Supplementary Data 3). For each comparison, a maximum of 1000 terms with an adjusted p-value < 0.15 from GSEA were included. Similarity between terms was calculated based on the Jaccard index, which quantifies the overlap between gene sets associated with each term. These similarity scores were used to cluster terms into functionally related groups. Each cluster was annotated using the term with the lowest adjusted p-value, representing the most significantly enriched term within the cluster. The summarized enrichment results (Supplementary Data 3) were visualized as heatmaps to facilitate comparison of biological pathway activity across conditions.

### Enrichment plots of specific gene sets and cell types

Plots showing GO terms enriched in DEGs identified in all NM pigmented areas (pan-regional) and SN and VTA-specific DEGS were obtained using ShinyGO website (http://bioinformatics.sdstate.edu/go/; RRID:SCR_019213). Cell-type–specific gene-ranking scores were obtained from CellKB (https://www.cellkb.com/; RRID:SCR_025985) for each gene across distinct single-cell datasets derived from the publicly available mouse brain atlas published by Saunders et al. (PubMed ID: 30096299)^30^.

### Correlation analysis (ROAST)

ROAST is a gene set test that allows for gene-wise correlation between two DEA ^31^. ROAST was performed using R software and the implemented function in Bioconductor Project (RRID:SCR_006442). This test was used to compare DEA in the tgNM model (SN region, LC region and SN neurons at different ages) to published studies characterizing transcriptomic profiles in human PD postmortem samples obtained from public database NCBI GEO Datasets. We retrieved 8 studies analyzing postmortem SN from PD and age-matched controls (GSE20163, GSE20164, GSE20292, GSE20333, GSE43490, GSE7621, GSE20141 and GSE24378) and one study analyzing postmortem LC in both conditions (GSE34516) from the public database NCBI GEO Datasets (RRID:SCR_005012). The alternative hypothesis considered in this statistical test depends on whether the DEGs in the dataset are expected to change in the same direction in the compared datasets or not, resulting in three different p-values representing three different alternative hypotheses tested: genes changing in the same direction in human as they do in mouse (up direction and down direction) and genes changing in mixed (up or down) directions. In our analysis, only mixed p-values are shown for simplification. Also, ROAST was used to compare the DEA performed in the tgNM model to 8 standardized PD-related gene sets from different databases, where no gene expression direction is assigned to the genes (ClinVar Gene-Phenotype Associations, DISEASES Curated Gene-Disease Assocation Evidence Scores, DISEASES Experimental Gene-Disease Assocation Evidence Scores, GAD Gene-Disease Associations, GWAS Catalog SNP-Phenotype Associations, GWASdb SNP-Disease Associations).

### GPNMB expression by quantitative PCR

RNA from microdissected tissue (mouse and human) and RNA from bulk postmortem human tissues was extracted using RNeasy Micro (#74004, Qiagen). Subsequent amplifation to cDNA was performed using Ovation® Pico WTA System V2 (#3302, Tecan Genomics), which amplifies cDNA from total RNA (500pg-50ng of total RNA). Amplification is initiated at the 3’ end as well as randomly throughout the RNA transcripts, enabling the amplification of degraded RNA. Specific primers for each validated transcript were designed using Primer-BLAST7, a National Center for Biotechnology Information (NCBI) tool for designing specific primers. Primers sequences are specified in Table 1. Primers were diluted in water and stored at −20°C at a stock concentration of 100µM. For gene expression analysis, only cDNA samples amplified from good quality RNA samples were used (RIN>5 or RIN>4 for human microdissected DA neurons). For qPCR, 10ng of cDNA from each sample in technical triplicates were mixed with the specific primers at a working concentration of 500nM and the 2X PowerUp™ SYBR™ Green Master Mix (#A25776, Applied Biosystem-ThermoFisher). qPCR was performed using the recommended cycling condition in a LightCycler® 480 System (Roche). Quantification analysis using Fit Points Method was performed with LightCycler Software and threshold cycles (CT) signals for each sample. Fold changes for each transcript and sample were calculated normalizing the arithmetic mean of technical replicates CTs to the geometric mean of three endogeneous control genes (Ndn, Rtn1, Ppia for mouse; FOXO1, UGGT1, GOLGA3 and HPRT1 for human bulk expression and HPRT1 and UGGT1 for human DA neurons expression) and then normalized to experimental control expression (wt for mice, Ctrl for human) using the comparative method (ΔΔCT method).

**Table 1.**
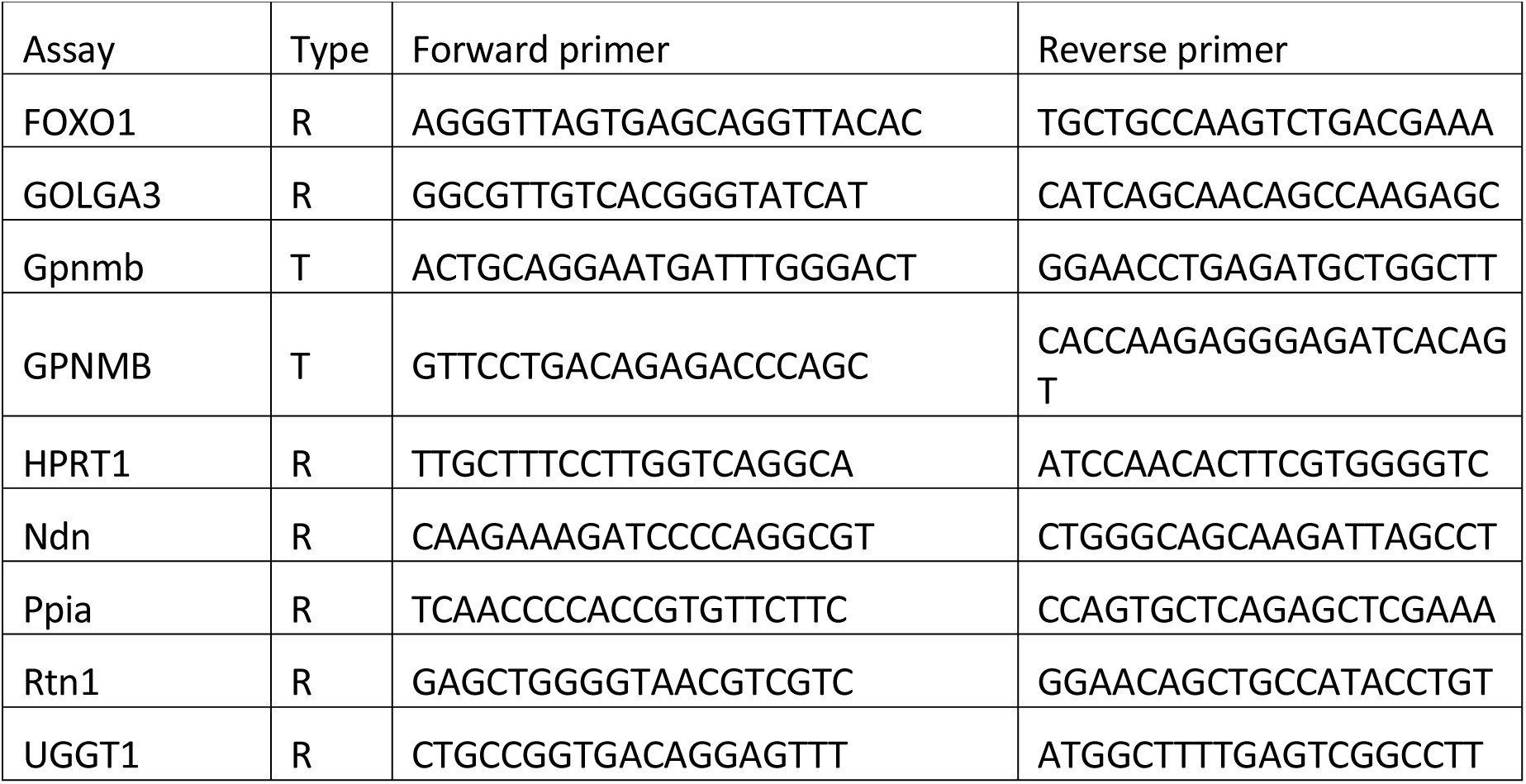
Primers designed using Primer-BLAST for mouse and human transcripts. Human transcripts are in capital letters. Target, T; Reference, R.

### Brain processing for histological analyses

Animals were deeply anesthetized with sodium pentobarbital (50 mg/kg, i.p.) and then perfused through the left ventricle with saline [0.9% (wt/vol)] at room temperature, followed by ice-cold formaldehyde solution 4% phosphate buffered for histology (Panreac). The brains were removed and post-fixed for 24 h in the same fixative and subsequently processed for paraffin embedding following standard procedures or cryoprotected for 24–48 h in 30% sucrose at 4 °C and frozen. Sectioning was performed in a cryostat at 20-thickness (CM3050S Leica, Germany). Sectioning of human postmortem brain regions was performed with a sliding microtome (RM2255 Leica, Germany) at 5 µm-thickness for paraffin samples.

### Immunohistochemistry and Immunofluorescence

Cryosections were quenched for 10 min in 3% H2O2-10% (vol/vol) methanol. Paraffin sections were deparaffinized and rehydrated. Sections were rinsed 3 times in 0.1 M Tris buffered saline (TBS) between each incubation period. Blocking for 1 h with 5-10% (vol/vol) normal goat, rabbit or donkey serum (Vector Laboratories, #S-1000 and #S-5000, and Sigma # D9663, respectively) was followed by incubation with the primary antibody (anti-TH, Calbiochem #657012, 1:40000 for SN and 1:5000 for striatum; Anti-GPNMB, R&D Systems-Biogen #AF2550,1:200 for human tissue; Anti-GPNMB, R&DSystems-Biogen #AF2330,1:1000 for mouse tissue; and Anti-GPNMB, Proteintech #66926, 1:500 for human tissue immunofluorescence; Anti-Iba1, Wako #019-19741;1:1000; Anti-Iba1, Abcam #ab178846;1:500 for immunofluorescence in human tissue) at 4°C for 24 or 48h in 2% (vol/vol) serum and with the corresponding biotinylated or Alexa Fluor antibodies (Vector Laboratories or Thermo Fisher Scientific, respectively) for 1h at room temperature. Sections were visualized directly or by incubation with avidin-biotin-peroxidase complex (Thermo Fisher Scientific, ABC Peroxidase Standard Staining Kit #32020 or Ultra-Sensitive ABC Peroxidase Standard Staining Kit #32050), using VectorSG Peroxidase Substrate Kit (Vector Laboratories, #SK-4700), DAB Peroxidase Substrate Kit (Vector Laboratories, #SK-4100) or ImmPACT Vector Red Substrate (Vector Laboratories, #SK-5105) as chromogens. Sections were then mounted and coverslipped with DPX or fluorescent mounting medium (Sigma-Aldrich 06522 or Dako S302380-2, respectively). Bright-field section images were examined with Zeiss Imager.D1 microscope coupled to an AxioCam MRc camera and with Pannoramic 250 Flash III (3D Histech), and processed with ZEN 2011 software (Zeiss, Germany, RRID:SCR_013672; https://www.zeiss.com/microscopy/en/products/software/zeiss-zen.html) and Caseviewer 3D HISTECH Ltd RRID:SCR_017654; https://www.3dhistech-com/caseviewer). For immunofluorescence analysis, maximum projection images (z-stack with a 0.20 μm interval) were taken in a confocal microscope (LSM 980; Zeiss) with AiryScan detection and an 40x oil immersion objective.

### RNAScope

Mouse paraffin sections were processed following the recommended protocol by Advanced Cell Diagnostics (ACD, Bio-Techne) RNAscope multiplex fluorescent reagent v2 assay (#323100). The RNAscope™ Probe used was Mm-Gpnmb (ACD, Bio-Techne #489511) together with TSA Vivid Fluorophore 650 (ACD, Bio-Techne #323273). RNAscope probe detection was combined with immunofluorescence for neuronal and microglial markers. Primary and secondary antibodies were: Anti-Iba 1 Abcam (ab178846; 1:500) and Anti-TH Chemicon (MAB5280; 1:1000) with goat anti-rabbit Alexa Fluor 488 (Thermo Scientific # A-11008, 1:1000) and goat anti-mouse Alexa Fluor 594 (Thermo Scientific # A-11005, 1:1000), respectively. To quench autofluorescence, TrueBlack® Lipofuscin Autofluorescence Quencher (20X in DMF) (Biotium; Cat#23007) was applied for 90 s after the final step (DAPI application) of the RNAscope assay procedure, and gently rinsed with PBS prior to coverslipping and imaging. Maximum projection images (z-stack with a 0.20µm interval) were taken in a confocal microscope (LSM 980; Zeiss) with AiryScan detection and an 40x oil immersion objective.

### Stereotaxic infusion of viral vectors

Recombinant AAV serotype 2/9 containing the human TYR cDNA driven by the cytomegalovirus (CMV) promoter (AAV-2/9-CMV-TYR; concentration 8,94 × 10^12^ gc/mL) or its empty vector (AAV-2/9-CMV-Null) were produced as previously described ^11^ at the Viral Vector Production Unit of the Autonomous University of Barcelona (UPV-UAB, Spain). Recombinant AAV vector serotype 2/9 expressing the human GPNMB cDNA driven by the cytomegalovirus (CMV) promoter (AAV2/9-CMV-h-GPNMB #AAV-210248, concentration 1 × 10^13^ gc/mL) was purchased from Vector Biolabs. For stereotaxic viral injection experiments, only male mice were used to reduce variability associated with sex-dependent hormonal cycling and to improve consistency in surgery-dependent outcomes and post-surgical behavioral phenotyping. Adult male C57BL/6J mice (Charles River; RRID:MGI:3028467), 10 weeks old at the time of surgery, were housed four per cage with ad libitum access to food and water during a 12-hour light/dark cycle. Surgical procedures were performed with the animals (42 male mice; 8 per group and time-post injection) placed under general anesthesia using isoflurane (3% for the induction phase and 1,5-2% for the maintenance phase) (Baxter). Vector solutions were injected using a 10 μL Hamilton Microliter Syringe fitted with a glass capillary (model 701, Cat#7635-01). Animals received 1 μL of a 1:1 mixture of AAV-GPNMB+AAV-EV, AAV-TYR+AAV-EV or AAV-TYR+AAV-GPNMB to achieve a final titration of 1 × 10^13^ gc/mL for AAV-GPNMB and 8,94 × 10^12^ gc/mL for AAV-TYR. In the TYR/EV and GPNMB/EV groups, viral vectors were diluted with vehicle (PBS-MK/40% Iodixanol). Injection was carried out unilaterally on the right side of the brain, right above the SN. Using a stereotaxic apparatus aligned to a flat skull position, the injection site was localized at the following coordinates relative to bregma, based on the stereotaxic Paxinos and Watson atlas: anteroposterior: −2.9 mm; medio-lateral: −1.3 mm; dorso-ventral: −4.2mm below dural surface. Infusion was performed at a rate of 0.4 μL/min and the needle was left in place for an additional 4 min period before it was slowly retracted.

### Stereological cell counting

At 2 months post-injection, the total number of SN TH-positive neurons, the number of SN NM-laden neurons and the total number of DA neurons in the SN of AAV-TYR and/or AAV-GPNMB injected mice were assessed. Micrographs of TH-immunostained SN serial brain sections were acquired with an Olympus Slideview VS200 slide scanner (RRID:SCR_024783). A specific artificial intelligence (AI)-assisted algorithm was implemented for the identification and quantification of TH-positive NM-negative neurons, TH-positive NM-positive neurons, TH-negative NM-positive neurons and eNM granules using the Olympus V200 Desktop 3.3 software (RRID:SCR_016167, https://www.olympus-lifescience.com/en/solutions-based-systems/vs200/). Serial 20-µm-thick paraffin-embedded sections covering the entire SN were included in the counting procedure (every sixth section, for a total of 10-11 sections analyzed/animal). The entire SN was considered in each section and each automated counting was manually revised by an investigator blinded to the experimental groups.

### Quantification of neuropathological parameters

The absolute number of extracellular NM aggregates was estimated within the same sections in which SN TH-positive stereological cell counts were performed (i.e. serial 20-µm-thick sections covering the entire SN, taking every sixth section, for a total of 10-11 sections analyzed/animal).

### Optical densitometry analyses

The density of TH-positive fibers in the striatum was measured by densitometry in serial coronal sections covering the whole region (4 sections/animal). TH-immunostained 20-µm-thick sections were scanned with an Olympus Slideview VS200 slide scanner (RRID:SCR_024783) and the resulting images were quantified using ImageJ software (NIH, USA; RRID:SCR_003070). Striatal densitometry values were corrected for non-specific background staining by subtracting densitometric values obtained from the cortex. Data are expressed as the percentage of the densitometric value of the equivalent anatomical area from the non-injected contralateral side of the same animal. Animals were analyzed at different time-points after AAV-TYR and AAV-GPNMB injection (n=6-8 per group). All quantifications were performed by an investigator blinded to the experimental groups.

### Cylinder behavioural test

To assess forelimb asymmetry, mice were individually placed in a transparent glass cylinder and recorded for a total of 5 minutes. Video recordings were subsequently analyzed to quantify forelimb use. The number of wall touches performed with the left and right forepaws was counted, and the percentage of contralateral (affected side) paw usage relative to total forelimb use was calculated. The cylinder was thoroughly cleaned with 70% ethanol between each session to eliminate olfactory cues. All behavioral testing was conducted during the light phase by an investigator blinded to the experimental groups.

### Statistical analysis

Statistical analyses were performed using GraphPad Prism v6 software (RRID:SCR_002798; http://www.graphpad.com/) using the appropriate statistical tests, as indicated for each figure legend. Since all experiments had a relatively small number of mice (n = 4–24 mice/group), nonparametric tests were used because a Gaussian distribution could not be assumed considering sample size. Depending on data distribution and group comparisons, either the Kruskal–Wallis test (for multiple group comparisons) or the Mann–Whitney U test (for two-group comparisons) was applied. Outlier values in qPCR experiments were identified by ROUT (Q = 1.0%) test and excluded from the analyses when applicable. All data are represented as box-and-whisker plots, showing the median, interquartile range, and individual data points.

## Results

### Age-dependent transcriptional profiling of anatomically defined pigmented brain areas

We previously reported^12^ that, in parallel to progressive age-dependent NM accumulation, tgNM mice exhibit noradrenergic neurodegeneration in the LC between 1 and 3 months (m) of age, followed by progressive dopaminergic dysfunction and incipient degeneration in the SN and VTA from 12 to 20 m (Fig. 1a-b). These pathological changes are accompanied by PD-like motor and non-motor alterations, αSyn Lewy-like pathology, immune activation, and secondary dysfunction/degeneration in interconnected non-catecholaminergic brain circuits (i.e., cholinergic and serotonergic), as observed in PD patients (Fig. 1b)^12^.

Given these progressive NM-linked alterations, we sought to characterize the underlying molecular events in pigmented catecholaminergic brain regions. We performed transcriptomic microarray analysis on anatomically defined, NM-pigmented areas selectively isolated by laser capture microdissection (LCM). Specifically, we microdissected the LC at 3 and 12 m, and SN and VTA at 3, 12, and 20 m of age (Fig. 1c), corresponding to different levels of age-dependent NM accumulation in these areas,^12^ along with the equivalent non-pigmented regions from age-matched wild-type (wt) animals for comparison (Fig. 1c). After an RNA quality control, the histological identification of the microdissected regions was validated by the expression of region-specific catecholaminergic markers (Suppl Fig. 1a-c).

Differential expression analysis (raw p-value <0.01 and FC>|1.5|) between pigmented (tgNM) and non-pigmented (wt) regions at different ages revealed an age-dependent pattern, with an increasing number of differentially expressed genes (DEGs) observed as NM accumulation progressed (Fig. 2a-c, Suppl Fig. 2a). Additionally, a regional-dependent pattern of differential expression was evident, with the LC exhibiting the highest number of DEGs, followed by the VTA and SN (Fig. 2d). The temporal and regional differences in the number of DEGs aligns with the degree of NM accumulation (Suppl Fig. 2b) and NM-linked pathology^12^ in these areas. In tgNM mice, LC neurodegeneration is already established by 3 m of age, while the SN and VTA exhibit dopaminergic dysfunction at later ages, with only incipient neurodegeneration.^12^ This is consistent with the higher number of DEGs observed in the LC compared to midbrain regions. Thus, the large number of DEGs in the LC likely reflects transcriptional changes occurring during and after neuronal degeneration, whereas DEGs in the SN and VTA may represent early dysfunctional transcriptional programs linked to intracellular NM buildup, potentially encompassing neuron-rescuing targets. The full list of DEGs for each region and age is available in our submitted data (Supplementary Data 1).

**Figure 2.**
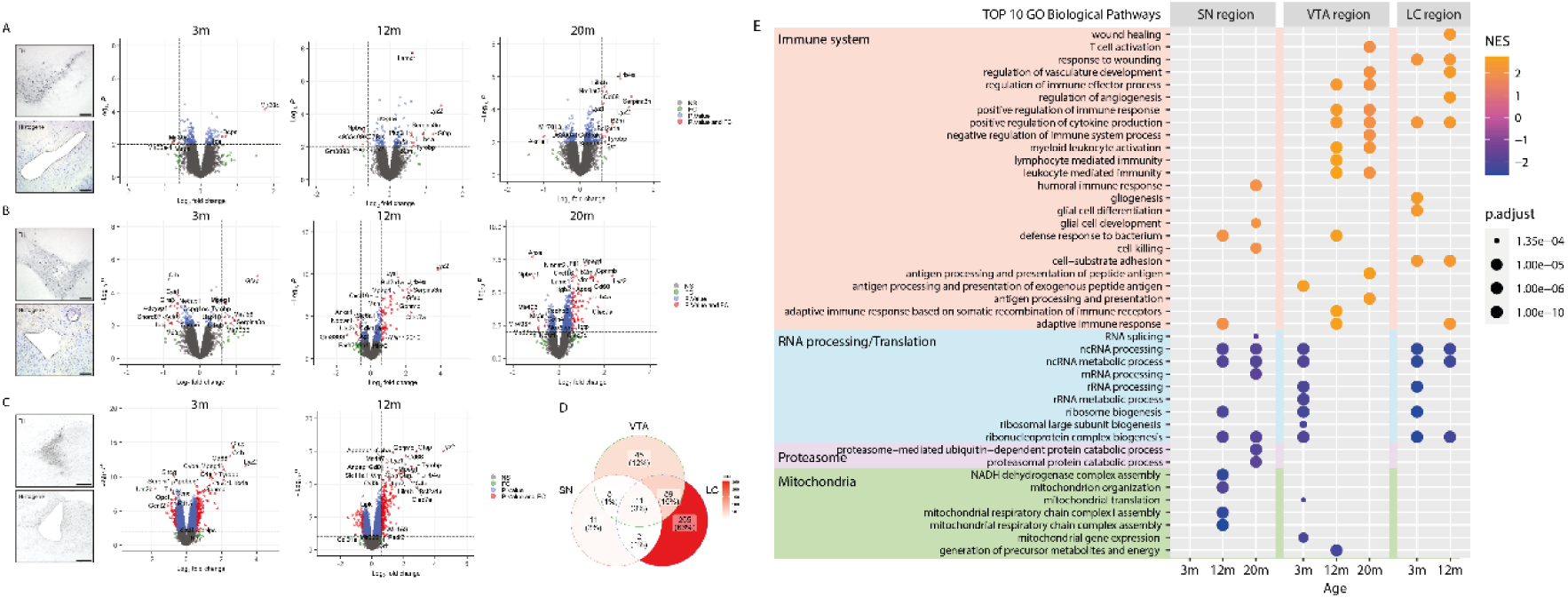
Region- and age-specific transcriptional changes associated with progressive neuromelanin accumulation in tgNM mice. **(a–c)** Volcano plots showing DEGs between tgNM and wt mice in the SN (a), VTA (b), and LC (c) at 3, 12, and 20 months of age. Adjacent panels display representative images of TH immunostaining and Histogene-stained sections used for LCM. Green dots indicate genes with fold change (FC > |1.5|), blue dots indicate genes with a raw p-value < 0.01, and red dots indicate genes that meet both criteria (p < 0.01 and FC > |1.5|), considered statistically significant DEGs. **(d)** Venn diagram summarizing the overlap of DEGs across SN, VTA, and LC regions at all ages. Color intensity reflects the number of genes in each region or overlap (count). **(e)** GSEA of up- and downregulated DEGs in SN and VTA neurons using the Gene Ontology (GO) annotation database. The top 10 significantly enriched biological pathways are displayed across the three catecholaminergic regions and ages. Dot color reflects normalized enrichment score (NES) and dot size corresponds to adjusted p-value (p.adjust). Enriched pathways are grouped into major biological categories: immune system (orange), RNA processing/translation (blue), proteasome (pink), and mitochondria (green).

Some DEGs were shared among the main pigmented regions (11 DEGs, 3%), while the majority were region-specific (Fig. 2d). We propose that these region-specific gene expression changes reflect the differential susceptibility of catecholaminergic populations to NM accumulation in tgNM mice, which exhibit earlier noradrenergic neurodegeneration in the LC compared to dopaminergic SN and VTA.^12^ Accordingly, we identified 11 SN-specific DEGs, 45 VTA-specific DEGs, and 255 LC-specific DEGs (Fig. 2d, Supplementary Data 1). The 11 common DEGs in all pigmented areas (i.e., B2m, Bcl2a1a, Cd68, Gfap, Lilr4b, Lilrb4a, Ly6a, Lyz1, Lyz2, Serpina3n, and Tyrobp) are predominantly expressed in inflammatory cells (macrophages/microglia and astrocytes) and associated with neuroinflammatory biological pathways, such as responses to external stimuli and cytolysis (Suppl Fig. 3a-b).

We also observed a clear enrichment of genes related to disease-associated microglia (DAM).^32–36^ Specifically, both Trem2 and its adaptor Tyrobp, a central hub of DAM signaling, were upregulated in all NM-containing areas from tgNM mice, supporting activation of the TREM2–TYROBP pathway. This pathway is a key regulator of microglial responses and is essential for the phagocytosis of apoptotic neurons^35,36^. To further investigate this enrichment, we performed a supervised search of established DAM-associated genes obtained from multiple studies characterizing microglial states in disease.^35–37^ This analysis revealed a consistent network of upregulated DAM genes present across all NM-accumulating regions (Suppl Fig. 3c, left panel). While Tyrobp showed robust, region-dependent upregulation, with the highest increases in the LC (FC ≈ 4.4-5.6, p ≤ 1.1×10⁻¹¹), intermediate changes in the VTA (FC ≈ 1.1–2.8, p ≤ 2.8×10⁻³) and weaker effects in the SN (FC ≈ 1.5, p ≈ 0.005), Trem2 upregulation was overall more modest and temporally delayed, with significant increases restricted to LC (FC ≈ 1.5, p ≤ 1.1×10⁻⁶) and VTA (12–20m: FC ≈ 1.2–1.4, p ≤ 0.003–2×10⁻⁵), while in SN it remained weak or non-significant across all ages (FC ≤ 1.2, p ≥ 0.005). Importantly, the expression of DAM-associated genes in the SN and VTA occurred concomitantly with dopaminergic dysfunction in these regions, despite the absence of overt neurodegeneration.^12^ As expected by the high degree of neurodegeneration in the LC,^12^ a marked upregulation of DAM genes was already detected at 3 m of age in this region. Additionally, we conducted a similar search for genes expressed in homeostatic microglia, and found differential expression predominantly in the VTA and LC, but not in the SN (Suppl Fig. 3c, right panel). This observation suggests that while in the VTA and LC there is a general increase in the microglial population together with a phenotypic display of DAM genes, in the SN, there is primarily an increase in specific DAM genes, indicating an early shift in the microglial phenotype. Together, these findings suggest region-specific differences in microglia-related transcriptional profiles, with patterns that may be consistent with an early shift in microglial phenotype in response to NM accumulation.

Like any other given cellular stimuli, NM accumulation may result in dependent co-regulated expression of complex gene sets that may not be reflected in single gene changes.^38^ To explore this, we identified functional gene sets enriched in pigmented areas compared to their equivalent non-pigmented counterparts. Gene set enrichment analysis (GSEA) performed using the Gene Ontology (GO) annotation database yielded statistically significant GO pathways in all regions and ages except in the SN at 3 m of age (Supplementary Data 2). Biological pathways (BP) related to neuroinflammation were found to be enriched in all three pigmented areas within the upregulated transcripts in tgNM mice (Fig. 2e). BP related to transcription and RNA processing/translation pathways were enriched within the downregulated genes in tgNM mice in all three areas, indicating a generalized downregulation of transcription activity linked to age-dependent NM accumulation in all catecholaminergic areas. Interestingly, within the top 10 altered BPs, proteasomal-related pathways appeared only in the SN, while mitochondrial-related pathways appeared in both the SN and VTA regions (Fig. 2e). Similar BP were obtained when GSEA was performed using the Reactome database (Supplementary Data 2). A redundancy reduction analysis based on the similarity between all BP identified in both annotation databases showed that neuroinflammatory pathways were among the most altered in all regions, and proteasomal function also appeared as one of the main alterations in the SN region (Suppl Fig. 4, Supplementary Data 3).

### Age-dependent transcriptional profiling of pigmented neurons

Our transcriptional characterization of pigmented regions revealed bulk alterations involving distinct cell types, with neuroinflammatory changes being one of the main consequences of NM accumulation in tgNM pigmented brain areas. To identify the neuron-specific effects of intracellular NM accumulation, we next aimed to characterize the neuronal transcriptome specifically in NM-filled neurons. This approach also aims to discern whether the neuroinflammatory process arises from an immune response to NM granules released into the extracellular milieu from dying neurons or from direct neuron-glia communication occurring prior to neuronal dysfunction and/or degeneration. To address these questions, we selectively isolated neuronal populations using LCM to better define the neuronal consequences of NM accumulation.

Given the extent of neurodegeneration observed in the LC region at early ages,^12^ we collected microdissected single neurons only from the SN and VTA (Fig. 3a, Video 1). Following microdissection and RNA quality controls (Suppl Fig. 5a), transcriptomic analyses confirmed the specificity of our approach by detecting established SN- and VTA-enriched neuronal transcripts (Suppl Fig. 5b). We detected a higher number of DEGs between tgNM and wt animals in neuron-enriched samples compared to regionally collected samples (Suppl Fig. 6a, Supplementary Data 1), supporting that NM accumulation primarily drives transcriptional changes in pigmented neurons, rather than in surrounding cell types. The neuron-targeted approach revealed a distinct transcriptomic profile in the SN compared to regionally collected samples, with only 6 overlapping genes out of 29 DEGs in the SN region (21%), whereas the VTA showed a higher overlap between both approaches, with 57 overlapping genes out of 120 DEGs (48%) (Suppl Fig. 6a). We validated our neuronal isolation protocol by assessing the cell type annotation of the main DEGs (top 500) using the CellKb database, which compiles single-cell datasets for cell type identification, reporting a higher proportion of neuronal DEGs in neuron-enriched samples compared to regionally collected samples (Suppl Fig. 6b). SN and VTA neurons exhibited a similar number of DEGs, with slightly more detected in SN neurons, primarily due to a large peak in DEGs at 12 m (Fig. 3b-c). Notably, the VTA displays a distinctive immune-related profile (e.g., Lyz2, Tyrobp, Cd68, Ms4a7, Gpnmb, etc.) emerging at 12 m and persisting at 20 m. The full list of DEGs for each neuron-enriched sample and age is available in our submitted dataenrich (Supplementary Data 1).

**Figure 3.**
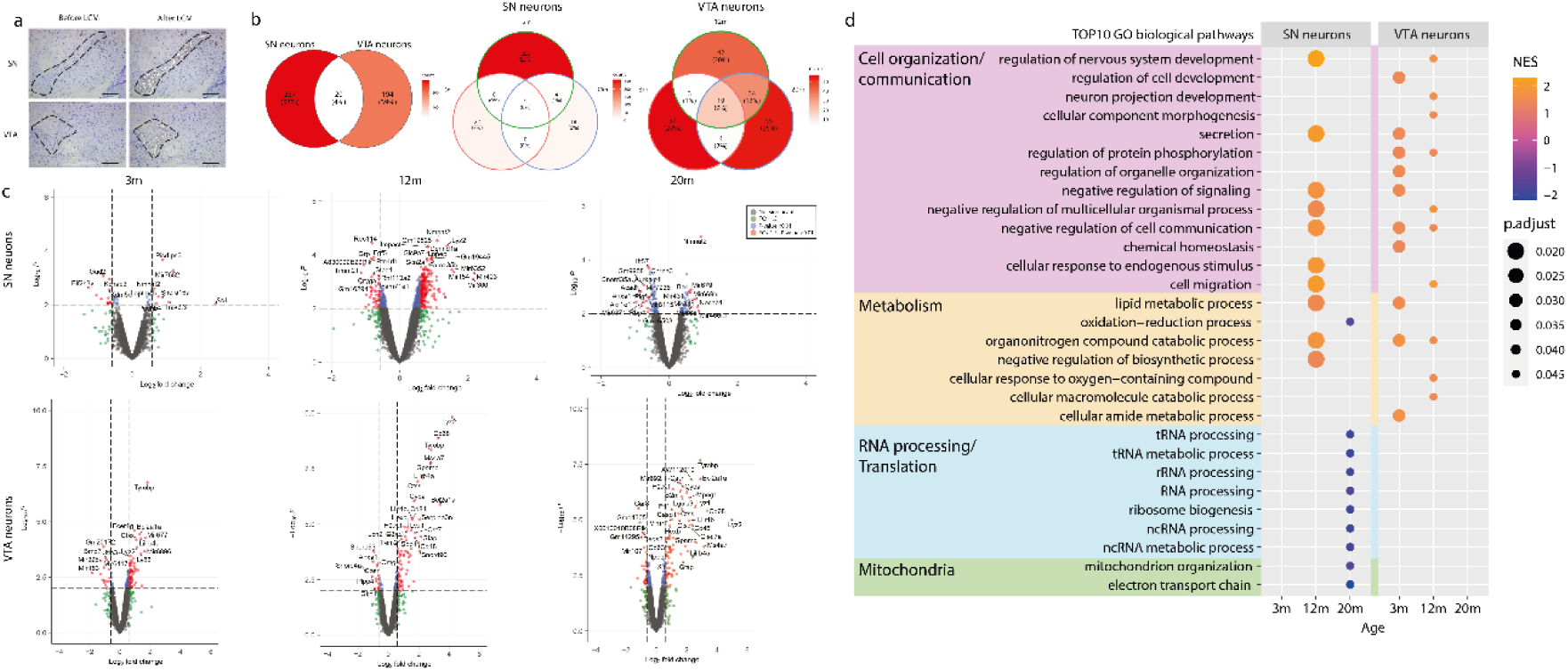
Neuron-specific transcriptomic alterations in pigmented dopaminergic neurons from tgNM mice. **(a)** Representative images showing LCM of pigmented neurons from SN and VTA, before and after microdissection. **(b)** Venn diagram summarizing the overlap of DEGs across SN and VTA neuron-enriched samples at all ages (left) and the overlap between DEGs identified at different ages (3, 12, and 20 months) in both regions (SN and VTA) (middle and right). Venn diagrams showing the overlap of DEGs between SN and VTA neuron-enriched samples at 3, 12, and 20 months, and their comparison with DEGs from regionally collected samples. Color scale represents the number of overlapping or DEGs per group.**(c)** Volcano plots of DEGs between tgNM and wt neuron-enriched samples in SN and VTA at 3, 12, and 20 months of age. Green dots represent genes with fold change (FC > |1.5|), blue dots represent genes with raw p-value < 0.01, and red dots indicate genes that meet both criteria and are considered significantly differentially expressed. **(d)** GSEA of up- and downregulated DEGs in SN and VTA neurons using the Gene Ontology (GO) annotation database. The top 10 significantly enriched biological pathways are displayed across the three catecholaminergic regions and ages. Dot color reflects normalized enrichment score (NES) and dot size corresponds to adjusted p-value (p.adjust). Enriched pathways are grouped into major biological categories: cell organization/communication (purple), metabolism (yellow), RNA processing/translation (blue), and mitochondria (green).

GSEA analysis using the GO annotation database revealed several distinct BP within the upregulated genes in the SN at 12 m of age, including those related to cell organization/communication and metabolism (Fig. 3d, Supplementary Data 2). Similar to regionally collected samples, pathways related to mitochondria and RNA processing were enriched among the downregulated genes, specifically in SN neurons at 20 m (Fig. 3d, Supplementary Data 2). Specifically, multiple mitochondrial pathways showed strong negative enrichment, including mitochondrial respiratory chain complex assembly (NES = −2.46, p = 0.0017), Complex I assembly (NES = −2.26, p = 0.0017), mitochondrial gene expression (NES = −2.25, p = 0.0018), mitochondrial translation (NES = −2.24, p = 0.0017), and ATP synthesis–coupled electron transport (NES = −2.11, p = 0.0018). In the previously analyzed regionally dissected SN samples, only the broader term mitochondrial organization was significantly enriched (NES = −1.56, p = 3.7 × 10⁻⁶). RNA processing pathways showed a similar pattern of partial concordance: 5 of the 40 RNA-related GO terms negatively enriched in LCM-isolated SN neurons and of the 13 RNA-related GO terms negatively enriched in regional SN samples, were significantly negatively enriched in both datasets (i.e., RNA methylation, RNA modification, ncRNA processing, ncRNA metabolic process, and tRNA metabolic process). These results support the notion that the dysfunctional phenotype observed in these pigmented dopaminergic regions at older ages may be driven by a generalized downregulation of neuronal transcriptional activity affecting various cellular functions. Conversely, VTA neurons exhibited a different pattern, with all enriched GO terms appearing among the upregulated genes and related to cell organization/communication and metabolism. No statistically significant biological pathways were enriched among the DEGs at 20 m of age in the VTA (Fig. 3b). Similar results were obtained when performing GSEA using the Reactome database. Redundancy reduction analysis of biological processes from both annotation databases highlighted pathways such as innate immune system, cellular signaling, neuronal system, membrane trafficking, and lipid metabolism (Suppl Fig. 7, Supplementary Data 3).

### Translational value of NM-induced changes in tgNM mice

To assess the similarity between the identified mouse NM-induced specific molecular fingerprints and human PD, we compared tgNM transcriptional profiles across multiple ages with publicly available datasets characterizing expression profiles in postmortem tissue from PD patients and age-matched controls. We retrieved eight studies analyzing postmortem SN (GSE20163, GSE20164, GSE20292, GSE20333, GSE43490, GSE7621, GSE20141, and GSE24378), as well as one study analyzing postmortem LC (GSE34516), from the public NCBI GEO database. No human transcriptomic studies specifically analyzing the VTA region were found in the literature. A detailed summary of these datasets is provided in our submitted data (Supplementary Data 4). Single-wise correlation analysis using ROAST ^31^ revealed a significant correlation of SN neurons at 20 m of age with four out of eight published human PD transcriptomic studies, and showed a clear trend toward correlation [p=0.05609 (GSE20141), p=0.05959 (GSE24378)] with two additional studies (Supplementary Table 1). Of note, only the transcriptomic profile of isolated SN neuronal populations from tgNM mice at 20 m of age showed correlation with postmortem PD SN, corresponding to the age with the highest levels of NM accumulation in tgNM mice. The sole study available analyzing LC from PD postmortem brains showed a clear correlation with the tgNM LC transcriptomic profile as early as 3 m of age, consistent with the early onset neurodegeneration reported in both tgNM mice and postmortem PD brains in this region (Supplementary Table 2). We then compared the same transcriptomic signatures to PD-associated gene sets from public databases based on experimental data. In this analysis, four out of six PD-associated gene sets showed a statistically significant correlation with tgNM SN neurons at 20 m, and all of them showed significant correlation with the LC transcriptomic signature (Supplementary Table 3-4). These results underscore the translational relevance of the tgNM model, as it recapitulates both region- and age-specific transcriptomic signatures observed in human PD.

### Validation of NM-induced GPNMB as a target in PD pathogenesis

We next proceeded to biologically validate the translational relevance of molecular candidates identified as NM-induced targets in PD pathogenesis. Among them, the melanosomal protein GPNMB (glycoprotein non-metastatic melanoma protein B) emerged as a promising candidate, as it was upregulated in all analyzed pigmented regions (i.e., SN, VTA, and LC) of tgNM mice, from young to old ages. GPNMB was initially identified as being expressed in melanocytes, where it is crucial for melanosome formation,^39,40^ but has since been shown to be expressed in various cell types and organs, including the brain, particularly in the context of neurological disorders.^41–43^ In PD, several studies have consistently shown GPNMB overexpression in postmortem SN tissue,^23,24,44–46^ and genome-wide association studies have linked the GPNMB locus to increased PD risk.^47–50^ Functionally, GPNMB has been implicated in macroautophagy,^51^ anti-inflammatory brain processes,^52^ the internalization of αSyn by neuronal cells^53,54^ and the clearing of intraneuronal αSyn mediated by microglia^55^. Despite the growing body of evidence, the exact role of this protein in the pathophysiology of PD remains unclear.

Gpnmb overexpression in pigmented brain areas of tgNM mice was confirmed by quantitative PCR (qPCR) (Fig. 4a). Gpnmb expression in these animals increased with age, paralleling the progressive accumulation of NM.^12^ In contrast, no Gpnmb expression was detected at any age in wt animals, which lack NM. Gpnmb expression also mirrored regional differences in NM levels, with earlier and higher Gpnmb overexpression in the LC, followed by the VTA and SN (Fig. 4a; Mean CT ± SD = 25.6 ± 1.8 [LC], 28.6 ± 7.4 [VTA], 35.9 ± 4.2 [SN]), reflecting the reported regional pattern of NM accumulation (LC > VTA > SN).^12^ Gpnmb overexpression was confirmed at the protein level by immunohistochemistry, showing co-localization of Gpnmb immunoreactivity with NM granules (Fig. 4b) and no detectable signal in non-pigmented wt animals (Suppl Fig. 8a). Combined RNAscope and immunofluorescence analyses revealed that Gpnmb expression was associated with both TH-positive neurons and microglial cells in tgNM mice (Fig. 4c and Suppl Fig. 8b). Collectively, these results confirm the identification of Gpnmb as an NM-dependent transcript whose expression is induced in both neurons and microglia in response to the progressive accumulation of NM in tgNM mice.

**Figure 4.**
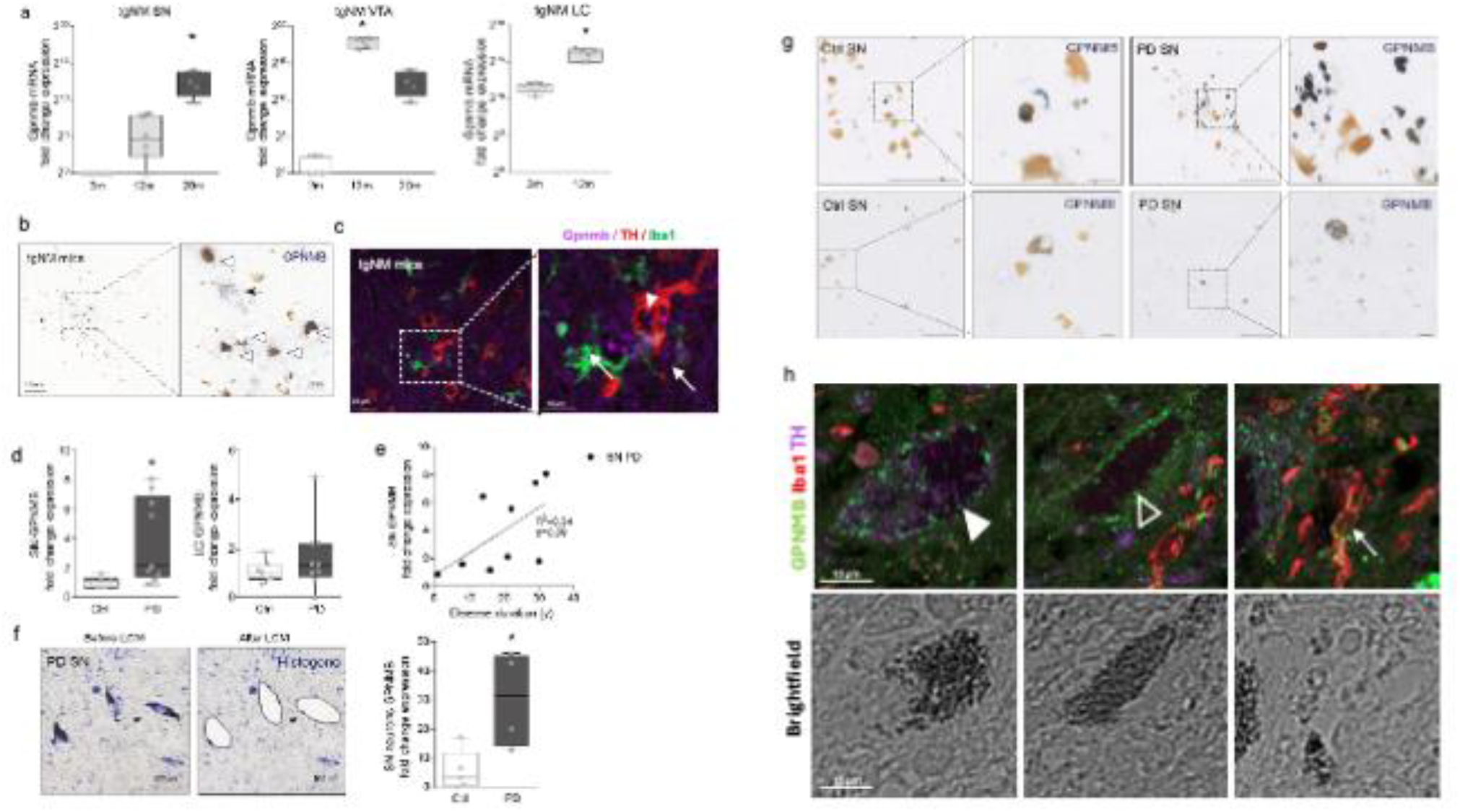
GPNMB expression in tgNM mice and PD postmortem brains. **(a)** Quantitative PCR analysis showing Gpnmb mRNA expression in neuromelanin (NM)-containing regions (microdissected SN neurons, VTA neurons and LC region) of tgNM mice at different ages. Levels of expression are calculated using the comparative method and shown in fold changes compared to wt and previously normalized to endogenous reference genes (Rtn1, Ppia, and Ndn). Almost no expression was detected in wt. *p≤0.05 compared to 3m; Kruskal Wallis and Dunn’s multiple comparison test (SN and VTA), Mann-Whitney test (LC). n=4-6 per group. **(b)** Representative image of Gpnmb immunohistochemistry in SN of tgNM mice at 20m of age. Gpnmb immunoreactivity is strongly expressed in tgNM mice, where it colocalizes with NM granules (brown pigment, open arrowheads) and sometimes staining neuronal shapes with faint NM (arrow). **(c)** Representative images of combined RNAscope and immunofluorescence staining in the SN of tgNM mice. Gpnmb transcripts (magenta) were detected in NM-containing neurons expressing tyrosine hydroxylase (TH, red; filled arrowheads). Gpnmb expression was also observed in Iba1-positive microglial cells (green; arrows) surrounding NM-containing neurons. Scale bar, 20 μm and 10 μm (High-magnification). **(d)** GPNMB mRNA expression in human postmortem SN and LC from PD patients and age-matched controls (Ctrl). Expression levels are calculated using the comparative method and shown as fold changes compared to control subjects and previously normalized with endogenous reference genes (FOXO1, GOLGA3, HPRT1, and UGGT1). *p≤0.05 compared to control (Ctrl), Mann-Whitney test. [n=6-7 (Ctrl), n=9 (PD)]. Samples with RIN<5 (SN Ctrl n=1, SN PD n=1, LC PD n=1) were excluded. **(e)** Linear regression analysis of GPNMB mRNA levels in PD SN samples (fold changes compared to Ctrl) as a function of disease duration in years post-diagnosis (years, y) (linear regression, p=0.09, R^2^=0.34). Dots represent individual samples. **(f)** LCM of pigmented neurons from PD SN, followed by qPCR analysis of GPNMB expression. Left, Histogene-stained sections before and after LCM illustrate the isolation of NM-containing neurons. Right, GPNMB expression levels in SN neurons from Ctrl and PD tissue expressed as fold changes compared to Ctrl previously normalized with endogenous control genes (HPRT1 and UGGT1). *p≤0.05 compared to Ctrl (Mann-Whitney test). [n=7 (Ctrl), n=7 (PD)]. Samples with RIN<4 (Ctrl n=2, PD n=2) were excluded. **(g)** Representative images of GPNMB immunohistochemistry in postmortem human SN from control and PD cases. GPNMB immunoreactivity primarily colocalizes with eNM granules but also sometimes localized within NM-containing cell bodies in both control cases (Upper left: 76 years Male; lower left: 83 years, Female) and PD (Upper right: Male, 77 years, iPD/PDD, Braak 5, 1 year of disease duration; Lower right: Male, 81 years, 29 years of disease duration. Scale bars: 200 µm (main images), 25 µm (insets). (**h**) Representative images of GPNMB double-immunofluorescence staining in postmortem human SN from PD cases (65 years, Male). GPNMB immunoreactivity is detected in TH-positive neurons with NM (filled arrowheads) and in TH-negative neurons with NM (open arrowheads), as well as in Iba1-positive microglial cells surrounding NM-containing neurons (arrows). Corresponding brightfield images highlight intracellular NM pigment. Scale bar: 10 µm

To assess the translational relevance of these findings to humans, we analyzed GPNMB expression in pigmented (SN, LC) and non-pigmented (frontal cortex, FC) regions from postmortem PD and control brains. Cohort characteristics, including clinical variables and experimental details, are provided in Supplementary Data 5, and GPNMB expression values in Supplementary Data 6. We first validated GPNMB RNA expression in cultured human melanocytes, a cell type in which GPNMB is highly expressed, using qPCR (Cycle Threshold [CT] mean ± SD = 18.8 ± 0.7), and subsequently confirmed its expression in human control brains, where expression levels were approximately 60 times lower (CT mean ± SD = 24.7 ± 1.6; Suppl Fig. 8c). In the SN, our analysis revealed an increase in GPNMB bulk RNA expression in PD brains compared to age-matched controls (Fig. 4d). Additionally, there was a trend toward a correlation between GPNMB levels in the SN and PD disease duration (linear regression, p=0.09, R²=0.34) (Fig. 4e). No statistically significant differences in GPNMB expression were detected between PD and control samples in the LC or FC (Fig. 4d, Suppl Fig. 8d). To further refine this analysis, we used LCM to isolate NM-containing neurons from SN of PD and control brains and confirmed increased GPNMB RNA expression in PD SN neurons (Fig. 4f; Video 2). Concordantly, GPNMB expression was confirmed at the protein level by immunohistochemistry in control and PD postmortem brains (Fig. 4g), showing high expression in clustered microglia as well as expression in large cells the size of dopaminergic neurons with NM. Double immunofluorescence analyses confirmed GPNMB expression in Iba1-microglial cells surrounding eNM as well as in NM-containing neurons (Fig. 4h and Supplementary Fig. 8e). Importantly, GPNMB-positive NM-containing neurons comprised both TH-positive and TH-negative populations. Given that downregulation of TH is a recognized feature of neuronal dysfunction in PD, these findings indicate that GPNMB expression is retained across dopaminergic neurons exhibiting different stages of pathological alteration. Overall, this distribution points to a potential dual role for GPNMB in both neuronal and microglial responses to NM-associated pathology.

To investigate the functional role of GPNMB in a NM-relevant PD context, we overexpressed GPNMB using an adeno-associated viral (AAV) vector in melanized nigral neurons of NM-producing mice. This approach involved AAV-mediated overexpression of human GPNMB or TYR, either individually or in combination, unilaterally in the SN of wt mice (Fig. 5a). Compared to tgNM mice, AAV-TYR-injected rodent models exhibit earlier and more pronounced NM accumulation, which is associated with accelerated nigrostriatal neurodegeneration.^11^ As expected, at 2 m post-injection (mpi), AAV-TYR-injected mice showed extensive ipsilateral nigrostriatal degeneration, characterized by loss of dopaminergic nigral cell bodies and nigrostriatal projections, along with motor asymmetry (Fig. 5b-c). Co-expression of GPNMB in these NM-producing animals provided a mild protective effect against dopaminergic neuron loss caused by NM accumulation, although no significant change was observed in striatal dopaminergic fibers. This protection correlated with improved motor performance in this PD-relevant model (Fig. 5c).

**Figure 5.**
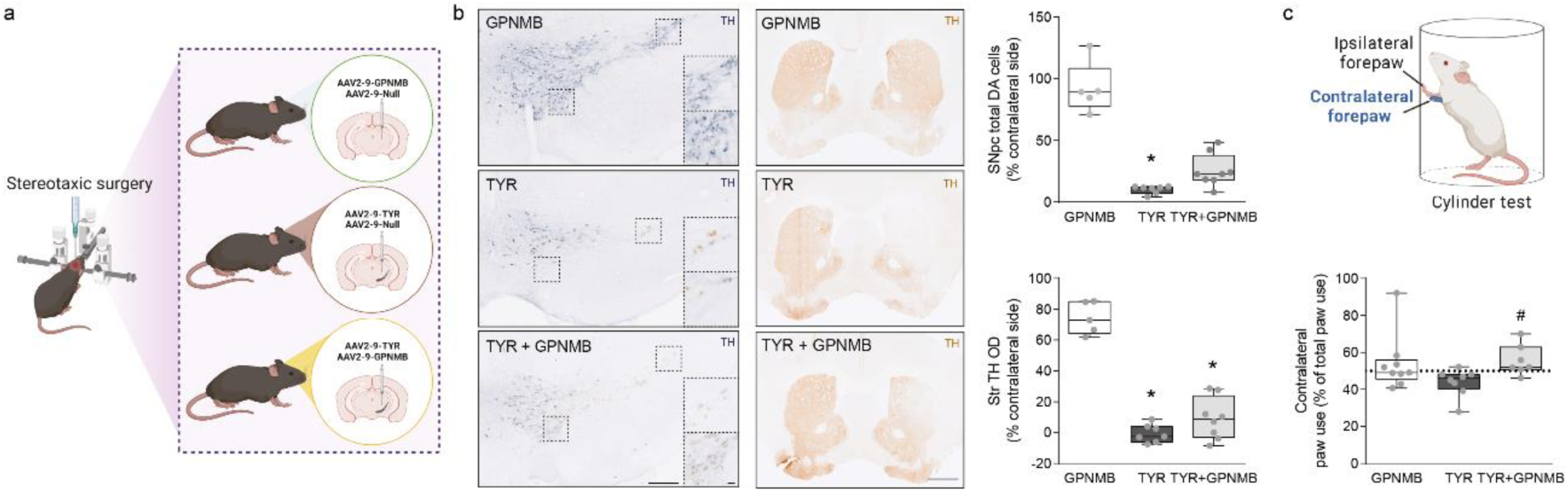
GPNMB overexpression attenuates TYR-induced dopaminergic neurodegeneration and motor deficits. **(a)** Schematic overview of the stereotaxic injection strategy. Mice received unilateral AAV injections into the SN to overexpress GPNMB alone (AAV2-9-GPNMB + AAV2-9-Null), TYR alone (AAV2-9-TYR + AAV2-9-Null), or both TYR and GPNMB (AAV2-9-TYR + AAV2-9-GPNMB). **(b)** Left: Representative TH-immunostained sections of the SN (scale bars: 200 µm; insets: 20 µm) and striatum (scale bar: 1,000 µm) from AAV-injected mice. Top right: Quantification of TH-positive and total dopaminergic (DA) neurons in the SN, expressed as a percentage of the contralateral (uninjected) side. Bottom right: Quantification of striatal TH optical density (OD), also expressed as a percentage of the contralateral side. *p ≤ 0.05 vs. AAV-GPNMB, Kruskal– Wallis test with Dunn’s multiple comparisons. **(c)** Schematic of the cylinder test used to assess motor asymmetry by quantifying contralateral versus ipsilateral forepaw use following unilateral AAV injection. Bottom: Quantification of contralateral forepaw use, expressed as a percentage of total paw contacts. #p ≤ 0.05 vs. AAV-TYR. In all panels, box plots represent median, minimum, and maximum values, with individual data points shown.

Overall, our transcriptomic analyses defined a molecular fingerprint of PD-vulnerable neurons associated with age-dependent NM accumulation, which correlated with gene expression profiles from postmortem PD studies and highlighted the disease associated gene GPNMB as a relevant target with potential therapeutic value.

## Discussion

This study provides a comprehensive investigation into the transcriptomic profiles associated with progressive NM accumulation in rodent SN, VTA, and LC, shedding light on the molecular mechanisms underlying PD-linked neuronal vulnerability of these pigmented brain regions.

Our study delineates NM-specific transcriptomic profiles across different pigmented brain regions, highlighting significant alterations in immune response, transcriptional regulation, protein synthesis, proteasome function, and mitochondrial activity. Notably, neuroinflammatory pathways emerged as a common feature across all pigmented areas, highlighting the important role of inflammation in NM-induced pathology. We further used LCM to isolate catecholaminergic neurons, providing insights into the neuronal consequences of NM accumulation. This approach revealed a higher number of DEGs in neuronal-enriched samples compared to regionally collected samples, emphasizing the importance of considering cell type-specific changes in NM-induced pathology. We detected fewer altered transcripts related to the immune system with this approach, meaning that in the regionally collected samples, transcriptomic changes may mainly represent glial reactions to released NM granules. Still, some immune transcripts were detected, mainly in VTA neurons. These changes may be attributed to direct neuron-glia contact, as LCM also captures the extracellular milieu surrounding collected neurons. Indeed, further studies using single-cell or single-nuclei techniques in NM-accumulating models may help elucidate such differences in cell-type specific changes.

The number of DEGs identified in tgNM mice pigmented areas followed a regional and age-dependent pattern similar to that of NM accumulation levels and neuropathological features, supporting the link of these changes to pigment accumulation. Main transcriptional changes in pigmented areas implicated known PD-related biological pathways (e.g. immune system, mitochondria, proteasome) and others that have not been so extensively linked with PD (e.g. transcription and translation, metabolism, cell communication), which might reveal novel therapeutic targets and pathways. Of note, we identified a common neuroinflammatory profile (B2m, Bcl2a1a, Cd68, Gfap, Lilr4b, Lilrb4a, Ly6a, Lyz1, Lyz2, Serpina3n, and Tyrobp) and a marked upregulation of distinct neuroinflammatory biological pathways in all pigmented areas in tgNM mice. This is consistent with the fact that neuroinflammatory changes are highly localized within NM accumulating areas in the human PD brain,^56,57^ and that some of these changes have also been reported in pigmented areas from non-diseased aged brains.^58,59^ Interestingly, the activation of immune pathways in tgNM mice was detected in SN and VTA regions, which exhibit dopaminergic dysfunction without overt neurodegeneration in these animals^12^. This finding suggests that the neuroinflammatory changes found in human PD and non-diseased aged pigmented brain areas^56–59^ may represent early pathological changes that precede overt neurodegeneration. Indeed, a number of PD-linked genes and risk factors have been identified as modulators of the immune function, postulating neuroinflammation as a potential key player in disease pathophysiology.^60^ The results presented here support the previously hypothesized role of NM in the initiation of a neuroinflammatory reaction in PD, either when accumulated inside NM-containing neurons and/or when released from dying neurons as eNM. We also detected an enrichment of the known DAM profile within pigmented areas. Importantly, previous immunohistochemical analyses in pigmented regions from the same tgNM model demonstrated an increase in reactive Iba1-positive microglia, while the number of non-reactive Iba1-positive cells remained unchanged, supporting a region-specific microglial activation state^12^. These reactive microglia were frequently associated with extracellular neuromelanin deposits, potentially reflecting phagocytic engagement with eNM released from degenerating neurons. These findings are consistent with the enrichment of DAM-related transcriptional signatures reported here. Together, these results might point to a microglial phenotype developed through the phagocytosis of eNM granules from dying neurons. However, further studies combining transcriptomic approaches (e.g., single-cell or single-nucleus RNA sequencing) with protein-level validation of DAM markers will be required to definitively confirm the existence of such specific phagocytic microglial phenotype able to engulf and degrade eNM granules.

One notable finding reported in this study is the correlation between the transcriptomic profiles identified in the tgNM model and human PD postmortem data. It should be noted that postmortem PD transcriptomes necessarily reflect a mixture of factors that could not be controlled for in our cross-species comparison, including pharmacological treatments, lifestyle, and aging. Still, the fact that a meaningful region- and age-specific correlation is observed, specifically in SN neurons at 20 months and in the LC at 3 months, suggests that NM-driven molecular changes in the tgNM model (assessed under controlled experimental conditions and in the absence of treatment- and lifestyle-related confounders) reflect a core transcriptional signature that can be found also in the heterogeneous background of the human disease. This observation further strengths the suitability of tgNM mice as a translational model for PD research and further implicates NM in the pathogenesis of PD. In the SN of the tgNM mice, the correlation with PD-derived data was only evident in the neuronal-enriched transcriptomic analyses and at 20m of age, thus suggesting that NM-related susceptibility to PD may be dependent on neuronal- and aging-related processes. The absence of significant correlation at earlier time points indicates that these stages may represent pre-symptomatic or early NM-associated molecular changes that have not yet progressed to a PD-like state. In contrast, a correlation of the tgNM mouse data with PD postmortem studies for the LC was already detected at 3 m of age, indicating that the reported extensive degeneration of the LC in young tgNM mice^12^ may already be equivalent to advanced stages of PD pathogenesis for this brain nucleus. Considering that NM-laden neuronal populations are also affected in other neurodegenerative diseases and aging, these findings extend beyond PD. Indeed, in control subjects, SN and LC neurons have been reported to decline with age,^61–68^ and in Alzheimer’s disease there is an early degeneration of the pigmented cells in the LC^69^ along with a reduced NM-occupied area in SN neurons.^70^ Hence, the molecular targets reported here may improve our understanding of how NM pigment accumulation contributes to catecholaminergic vulnerability not only in PD, but also to a variety of neurologic diseases and aging *per se*.

Additionally, we identified GPNMB as a key NM-induced target in the transcriptomic profiles presented here and validated its overexpression in both tgNM mice and human PD samples. Our findings of Gpnmb overexpression specifically in NM-accumulating tgNM animals align with *in vivo* studies in PD models, which show that GPNMB overexpression is only observed after lysosomal dysfunction, while synucleinopathy alone is insufficient to trigger this response^44^. This observation is particularly compelling given GPNMB’s involvement in melanosome formation and macroautophagic processes, suggesting a potential role in the formation and fusion of NM-filled autophagic vacuoles, ultimately leading to the development of NM-containing organelles^40,71^. This supports the notion that lysosomal stress in NM-laden neurons, caused by the progressive, age-dependent accumulation of NM within autophagic structures, as observed in humans^71^ and NM-accumulating *in vivo* models ^11^, may play a role in GPNMB overexpression. Furthermore, our results showing GPNMB upregulation in PD postmortem brains are consistent with previous studies showing consistent upregulation of GPNMB in human PD postmortem SN tissue and plasma^23,24,44,46^. We also report a positive correlation between GPNMB levels and disease duration, although this correlation was not observed in a prior study that found no association between GPNMB levels in the SN and disease duration, Braak stage, neuronal loss, or extraneuronal NM^44^. However, recent studies have linked GPNMB plasma and CSF levels to severity of motor and cognitive dysfunction^53,72,73^, suggesting its involvement in PD progression. An important unresolved issue is the precise cellular origin of GPNMB expression in pigmented brain regions. In the present study, we identify GPNMB-positive cells in NM-rich areas that represent both neurons undergoing phenotypic alterations associated with progressive NM accumulation and microglia engaged in phagocytosis of eNM released from degenerating neurons. Most studies suggest GPNMB is expressed by microglia, including recent single-nucleus RNA sequencing studies of postmortem human brain, which have identified GPNMB as a marker of a subset of microglial cells upregulated in PD and other neurodegenerative conditions^23,24,34^. However, other studies have shown GPNMB expression in multiple cell types, including neurons, in control and PD brains^34,45,53,54^. In agreement with this, we observed increased GPNMB expression in LCM-isolated NM-laden SN neurons from PD cases compared to controls, consistent with a similar observation in a previous study using the same approach^46^. We also provide evidence that GPNMB can attenuate NM-linked neurodegeneration *in vivo*. The protective effect of GPNMB may be linked to its ability to regulate the neuromelanogenic process or to other known functions, such as its anti-inflammatory properties^52,74,75^. A protective role for GPNMB has also been described in a toxic PD model^76^, though not in synucleinopathy-based PD models^45^, where GPNMB overexpression has no observed effect. These discrepancies may be due to differences in the PD subtypes modeled, with synucleinopathy-based models potentially representing a distinct PD subtype. It should be noted that the current study demonstrates that neuronal overexpression of GPNMB is sufficient to confer partial neuroprotection in a TYR-driven NM-producing model, while whether endogenous GPNMB is necessary for this protection remains to be established through future loss-of-function studies. Overall, the results of this study highlight a potential therapeutic role of GPNMB in PD and in the aging-related degeneration of pigmented neuronal populations.

## Conclusion

This study provides a comprehensive characterization of the molecular consequences of age-dependent NM accumulation in catecholaminergic brain regions using the NM-producing tgNM mouse model. Our findings demonstrate that NM accumulation drives region- and age-specific transcriptional changes, including the upregulation of neuroinflammatory pathways, alterations in transcription, translation, proteasomal and mitochondrial function, and modulation of neuronal metabolism and signaling. Laser capture microdissection of neurons revealed that many of these changes are neuron-intrinsic, rather than solely glial-mediated, highlighting the direct impact of NM on neuronal physiology.

Importantly, the transcriptomic profiles of NM-laden neurons in tgNM mice correlate with human postmortem PD data, underscoring the translational relevance of this model. The identification and validation of GPNMB as a key NM-induced target, with expression in both tgNM mice and human pigmented neurons, together with functional studies showing that it can mitigate NM-linked neurodegeneration *in vivo*, further demonstrates the potential of NM-related pathways as translational therapeutic targets.

Overall, our results provide strong evidence that NM accumulation contributes to neuronal vulnerability in PD by inducing specific transcriptional programs and engaging immune-related pathways. Beyond PD, these findings shed light on aging-related degeneration of pigmented neurons, offering a molecular framework to understand how age-dependent NM accumulation affects neuronal function and resilience. These insights pave the way for the development of targeted interventions and biomarkers for NM-associated neurodegenerative processes and age-related neuronal decline.

## Video legends

**Video 1. Laser capture microdissection (LCM) of single neuromelanin (NM)-containing neurons from the substantia nigra (SN) in tgNM mice.** The video illustrates the isolation of individual NM-containing neurons from anatomically defined pigmented SN in tgNM mice using LCM.

**Video 2. Laser capture microdissection (LCM) of neuromelanin (NM)-containing neurons from postmortem substantia nigra (SN) of an aged control brain.** The video illustrates the isolation of individual NM-containing neurons from human SN tissue using LCM.

## Abbreviations

AAV: Adeno-associated viral vector
BP: Biological pathway
CT: Cycle threshold
DA: Dopaminergic
DEA: Differential expression analysis
DEG: Differentially expressed gene
eNM: Extracellular neuromelanin
FC: Frontal cortex
GPNMB: Glycoprotein non-metastatic melanoma protein B
GSEA: Gene set enrichment analysis
LC: Locus coeruleus
LB: Lewy body
LCM: Laser capture microdissection
m: Months (age of mice)
NM: Neuromelanin
PD: Parkinson’s disease
qPCR: Quantitative polymerase chain reaction
RIN: RNA integrity number
RNA: Ribonucleic acid
SN: Substantia nigra pars compacta
TH: Tyrosine hydroxylase
tgNM: Transgenic neuromelanin mouse (Tyrosinase-overexpressing mouse)
TYR: Tyrosinase
VTA: Ventral tegmental area
wt: Wild-type

## Declarations

### Ethics approval and consent to participate

All procedures involving human samples were conducted in accordance with the guidelines established by the Good Clinical Practice, GCP (CPMP/ICH/135/95) and Spanish regulation (223/2004), and were approved by the Vall d’Hebron Research Institute (VHIR) Ethical Clinical Investigation Committee (PR(AG)370/2014). Written informed consent was obtained from all participants or their legal representatives, in compliance with applicable regulations.

All experimental and surgical procedures involving animals were performed in strict accordance with European (Directive 2010/63/UE) and Spanish laws and regulations (Real Decreto 53/2013; Generalitat de Catalunya Decret 214/97) on the protection of animals used for experimental and other scientific purposes. Animal protocols were approved by the Vall d’Hebron Research Institute (VHIR) Ethical Experimentation Committee and the Generalitat de Catalunya (Protocol 11442).

### Consent for publication

Not applicable

### Availability of data and materials

All raw data generated in this study are provided in the Supplementary Information/Source Data File and are available in the Zenodo database under accession code https://doi.org/10.5281/zenodo.21719879. All detailed protocols have been uploaded in the públic repositiry Protocols.io under the accession code: https://www.protocols.io/view/age-dependent-brain-pigmentation-drives-early-neur-2bvjro11plk5/v1. Further information and requests for resources and reagents should be directed to and will be fullfilled by the Lead Contact, Miquel Vila.

### Clinical trial number

Not applicable.

### Author contributions

Conceptualization and supervision: A. Laguna, M. Vila

Experimental design, planning, and oversight of all aspects of the study: N. Peñuelas, A. Laguna, M. Vila

Acquisition of experimental data and analysis: N. Peñuelas, H. Xicoy, M. Lorente-Picón, A. Nicolau, A. Parent

Data curation and management: J. Hoyo

Manuscript drafting: N. Peñuelas, A. Laguna, M. Vila, with input from all authors

All authors read and approved the final manuscript.

## Acknowledgements

We are grateful to: the Neurological Tissue Bank of the Biobanc-Hospital Clinic-IDIBAPS and the Vall d’Hebron Hospital Neurological Tissue Bank (Barcelona, Spain) and the Biobanco en Red de la Región de Murcia-BIOBANC-MUR (Murcia, Spain) for human sample procurement; Guillem Colell from the Laboratory Animal Service at VHIR, Dr. Takafumi Hasegawa (Department of Neurology, Tohoku University School of Medicine, Sendai, Japan) for kindly providing the TYR plasmid; to Pilar Mancera Aroca and Maria Molinos Marquez from Unitat d’Alta Tecnologia (UAT) at VHIR for their expert help with microarray analyses and dedicated suport throughout the project; to Ricardo Gonzalo Sanz and Mireia Ferrer from the Bioinformatics Platform of the Statistics and Bioinformatics Unit (UEB) at VHIR for their insightful bioinformatic support. All diagrams and schematic experimental plans were created with BioRender.com. For the purpose of open access, the author has applied a CC-BY 4.0 public copyright license to all Author Accepted Manuscripts arising from this submission.

