## Supplementary Material for "Age-dependent brain pigmentation drives early neuroinflammatory molecular signatures linked to neurodegeneration"

### Supplementary Materials

**Supplementary Figure 1. Quality control of laser capture microdissection (LCM)-isolated brain regions and transcriptomic profiling.** (a) RNA integrity number (RIN) values of LCM-isolated samples plotted as a function of dissection time, showing consistently high-quality RNA across samples. The dashed line indicates a commonly used RIN threshold of 5. (b) RIN values (left) and RNA concentrations (pg/μL; right) of neuromelanin-pigmented SN, VTA, and LC regions isolated from wild-type (wt) and tgNM mice. Across all brain regions and genotypes, samples showed acceptable RNA quality and yield suitable for microarray analysis. (c) Validation of anatomical specificity of LCM-isolated regions by assessing the expression levels of region-enriched transcripts in wt animals. SN markers (Aldh1a1, Pitx3, Kcnj6) were highly enriched in the SN compared to VTA and LC regions; VTA markers (Otx2, Calb1, Neurod6) showed selective expression in VTA samples; and LC markers (Dbh, Phox2a, Phox2b) were enriched in LC samples. Expression values represent raw microarray expression levels. *p<0.05 Two-way ANOVA. (a-c) Boxplots represent median and interquartile range with individual data points.

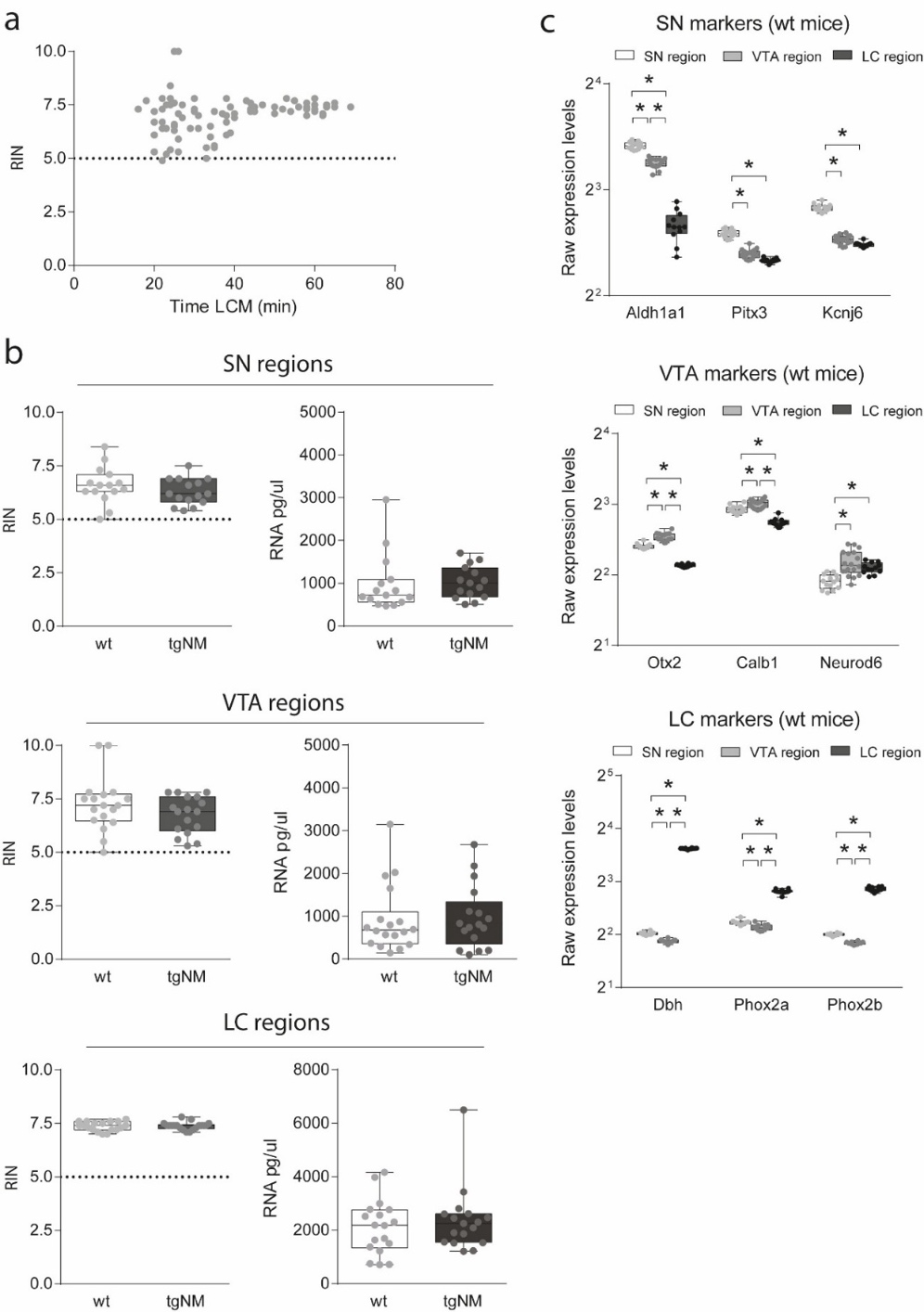
**Supplementary Figure 2. Age-dependent dynamics of differentially expressed genes (DEGs) in neuromelanin (NM)-pigmented brain regions.** (a) Venn diagrams showing the number and overlap of DEGs in the substantia nigra (SN), ventral tegmental area (VTA), and locus coeruleus (LC) at different ages (3, 12, and 20 months). Color scale represents the number of genes in that particular region or overlap (count). In the SN and VTA, a progressive increase in unique and shared DEGs is observed with age, peaking at 20 months. In contrast, the LC shows a large number of DEGs already at 3 months, with continued accumulation at 12 months, consistent with earlier onset of neurodegeneration in this region.
(b) Relation between the number of DEGs and mean optical density (OD) of intracellular NM in SN (circles), VTA (squares), and LC (triangles) across ages quantified in independent animals. A positive relationship is observed between NM accumulation and transcriptional dysregulation, supporting the association between pigment deposition and molecular pathology in catecholaminergic neurons.

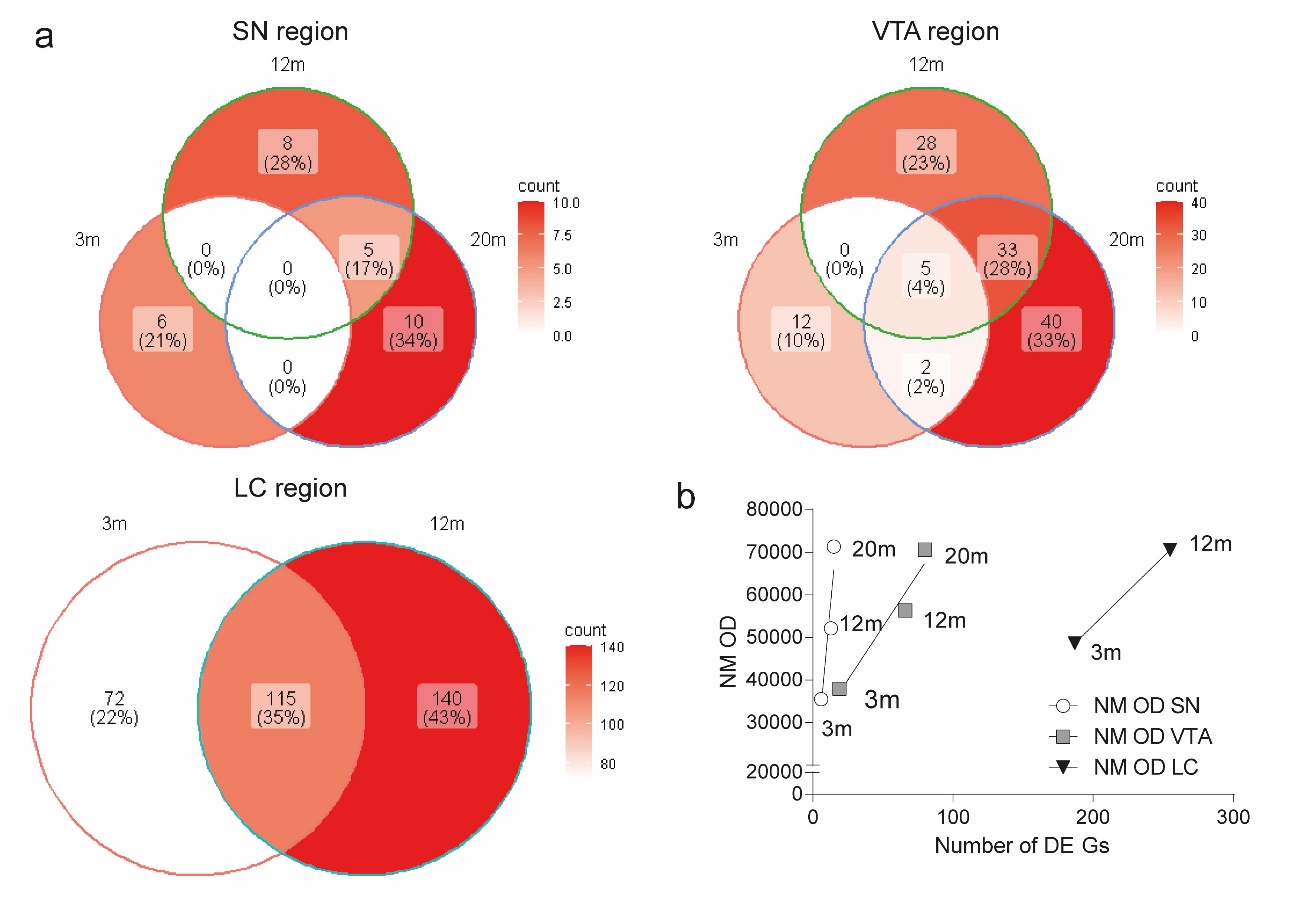

**Supplementary Figure 3. Pan-regional neuromelanin (NM)-linked transcriptional changes across catecholaminergic brain regions.** (a) Differentially expressed genes (DEGs) across all three NM-pigmented regions (LC, SN, and VTA) are predominantly associated with a neuroinflammatory profile. These include transcripts specific to the myeloid lineage (e.g., microglia/macrophages), astrocytes, and to a lesser extent, endothelial and neuronal populations. The cell-type enrichment for each gene was assessed using gene-rank scores from the CellKB platform (<https://www.cellkb.com/>) based on the single-cell transcriptomic reference dataset from Saunders et al. (PMID: 30096299)(Saunders et al., 2018). (b) Gene ontology (GO) enrichment analysis of the 11 DEGs shared by all NM-pigmented regions was performed using ShinyGO. Shown are the significantly enriched GO terms, along with the number of overlapping genes, fold enrichment, and false discovery rate (FDR). (c) DEGs (p ≤ 0.01) related to microglial activation states across all samples analyzed. On the left, transcripts characteristic of homeostatic microglia; on the right, transcripts known to be upregulated in microglia during the phenoconversion to DAM extracted from ^14,16,20^. Genes with positive log fold change (logFC) are shown in red, indicating upregulation; those with negative logFC are shown in light blue, indicating downregulation.

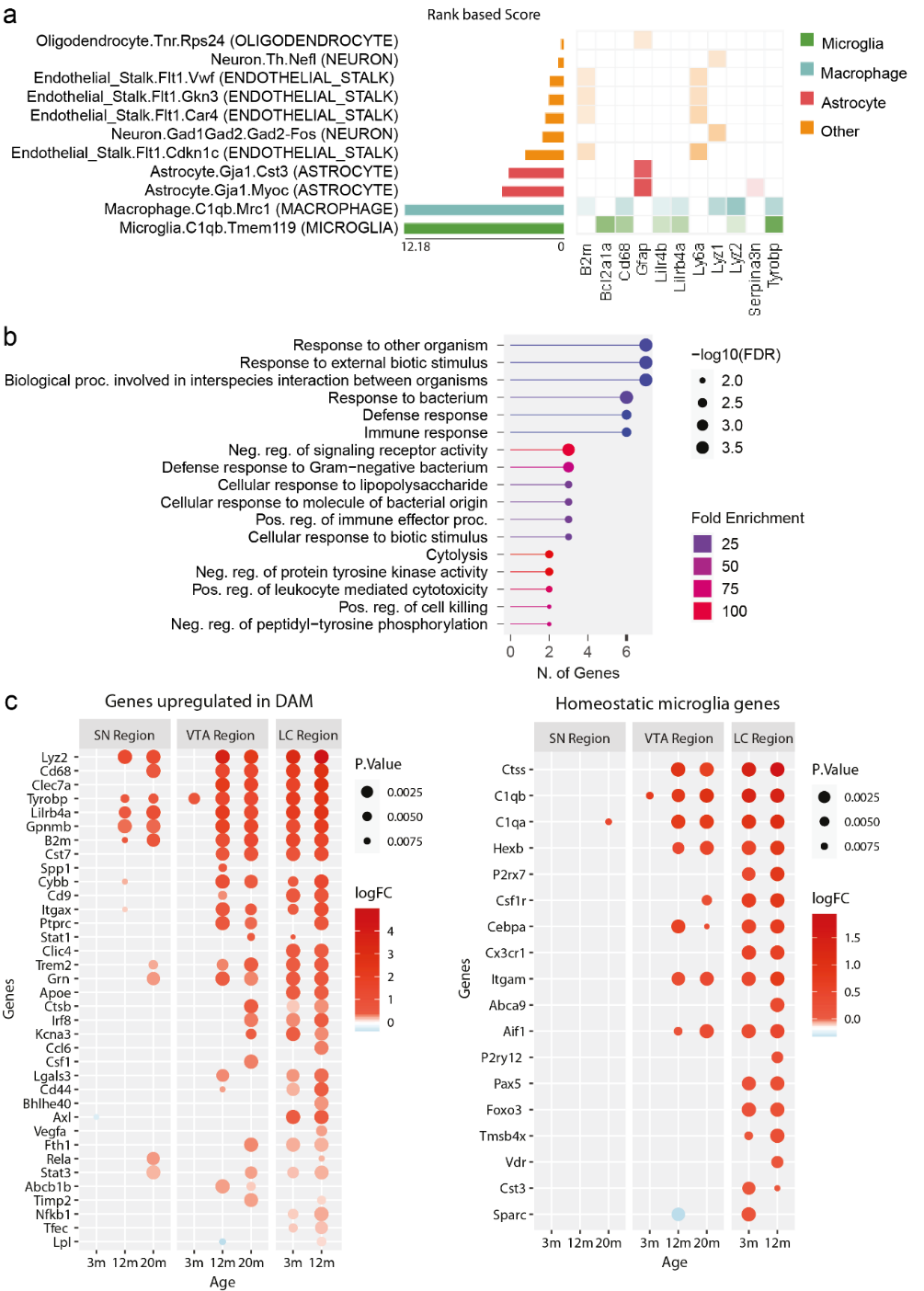

**Supplementary Figure 4. Gene-set enrichment analysis (GSEA) of biological pathways in neuromelanin-pigmented brain regions.** Similarity between enriched pathways from the Gene Ontology Biological Process (GO BP) and Reactome databases is depicted using the Jaccard Index for the substantia nigra (SN), ventral tegmental area (VTA), and locus coeruleus (LC) of tgNM mice relative to wild-type (wt) controls across different ages (3, 12, and 20 months). Pathways were identified by GSEA based on the Normalized Enrichment Score (NES), reflecting the degree of coordinated transcriptional changes associated with neuromelanin accumulation and neurodegeneration. The Jaccard Index quantifies the overlap between pathway gene sets, synthesizing biological processes underlying age-dependent molecular alterations in these catecholaminergic nuclei.

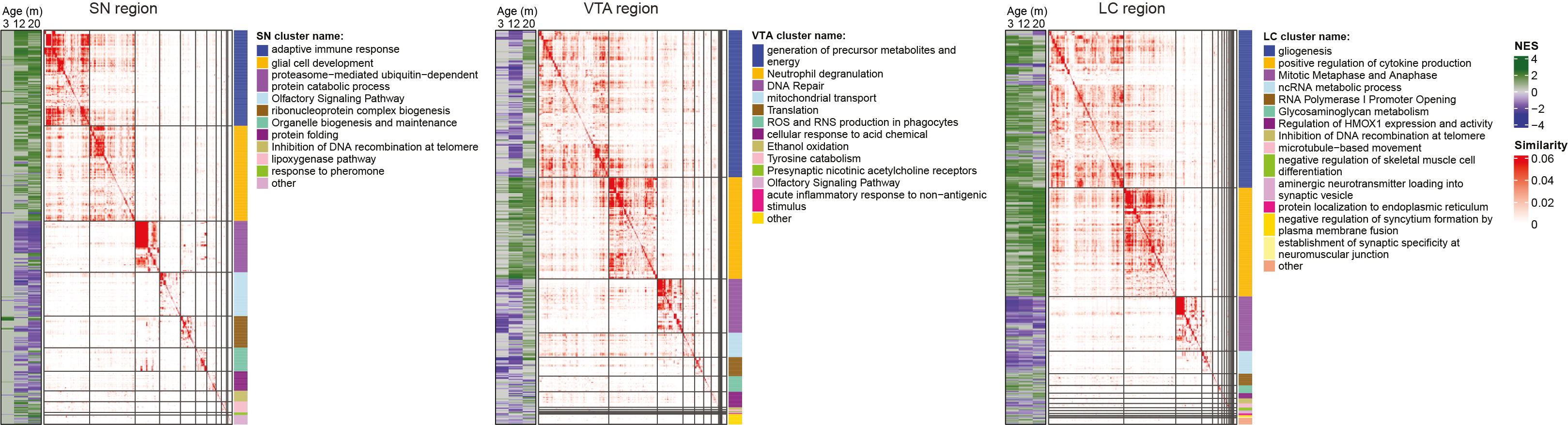

**Supplementary Figure 5. Quality control of LCM-isolated dopaminergic neurons and validation of neuronal subtype identity.** (a) RNA integrity number (RIN, left) and RNA concentration (pg/μL, right) of laser-captured neurons from the substantia nigra (SN) and ventral tegmental area (VTA) of wild-type (wt) and tgNM mice. Samples across genotypes and ages exhibited RNA quality and yield suitable for transcriptomic analysis. (b) Expression levels of neuronal subtype-defining markers in SN and VTA neurons isolated from wt animals across all ages. SN-enriched genes (e.g., Aldh1a1, Slc6a3, Sox6) and VTA-enriched genes (e.g., Calb1, Otx2, Cck) display distinct expression patterns consistent with dopaminergic subtype identity. Expression values represent raw microarray expression levels. *p ≤ 0.05 (Mann–Whitney test). (a-b) Boxplots represent median and interquartile range with individual data points.

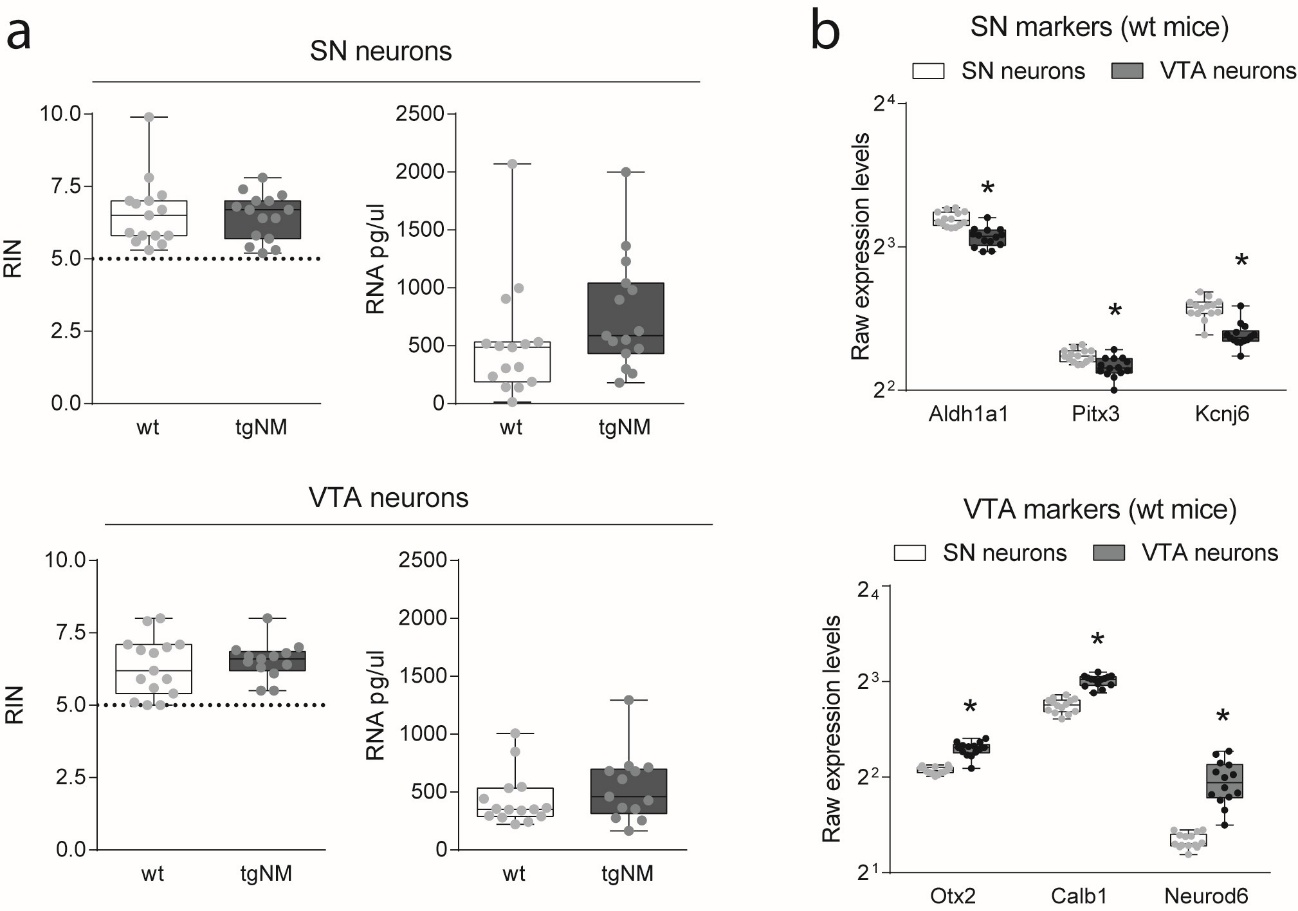

**Supplementary Figure 6. Comparison of differentially expressed genes (DEGs) identified from neuromelanin-pigmented regions versus isolated neurons.** (a) Overlap between DEGs identified in the substantia nigra (SN) and ventral tegmental area (VTA) of tgNM mice compared to non-pigmented wild-type (wt) controls, using two distinct LCM strategies: whole-region collection versus neuron-specific isolation. The red color gradient indicates the number of overlapping DEGs (count) between the two methods. (b) Cell-type annotation of the top 200 DEGs from SN regions, SN neurons, VTA regions and VTA enurons at 20 months, using gene enrichment scores from the mouse single-cell reference dataset by Saunders et al. (2018, PMID: 30096299) via the CellKb platform (<https://www.cellkb.com/>). Neuronal genes are shown in green, while non-neuronal clusters (e.g., glial and endothelial cell markers) are shown in orange, illustrating the cellular composition of DEGs captured by each method.

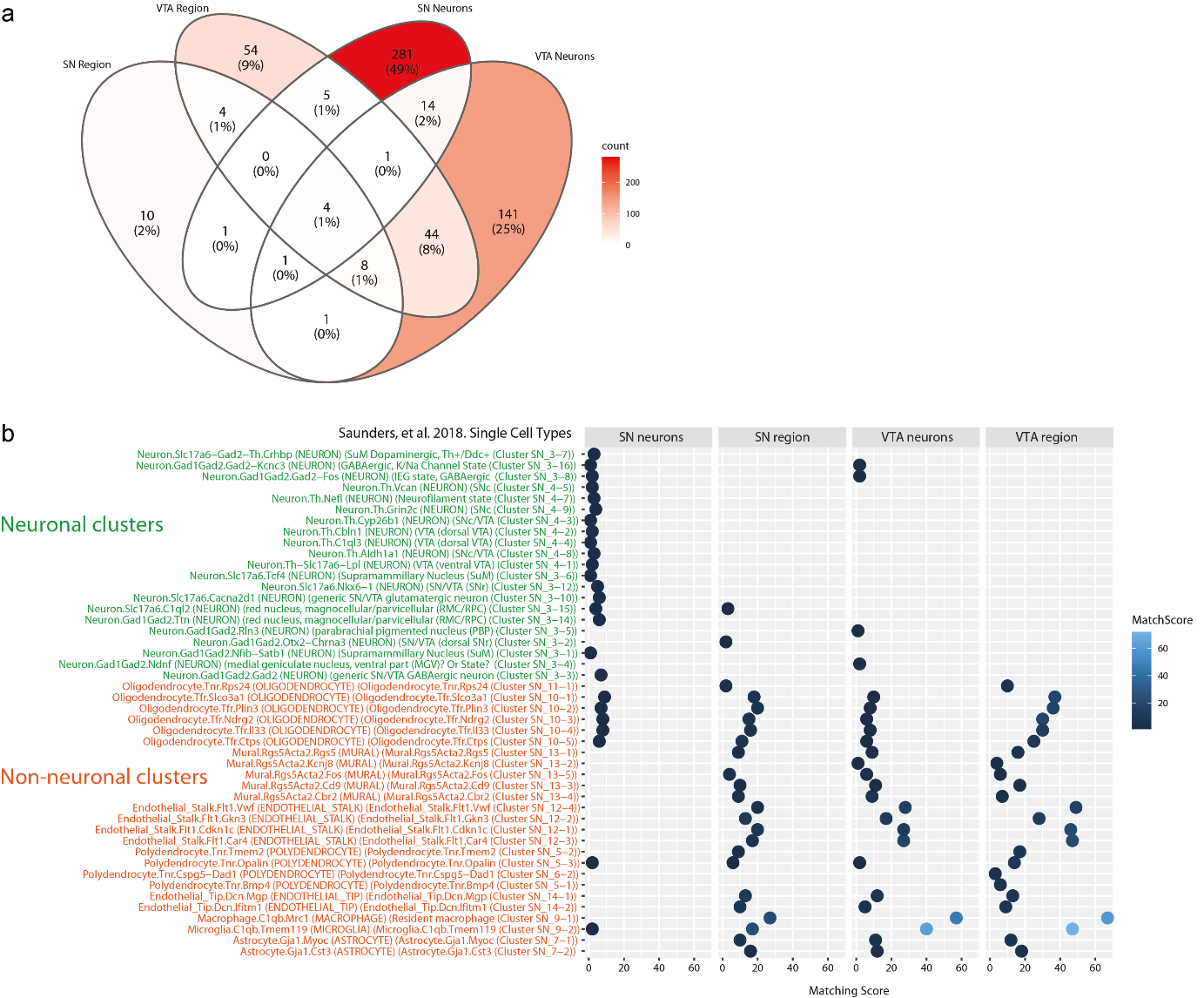

**Supplementary Figure 7. Gene-set enrichment analysis (GSEA) of biological pathways in neuromelanin-pigmented dopaminergic neurons.** Similarity between enriched pathways from the Gene Ontology Biological Process (GO BP) and Reactome databases is depicted using the Jaccard Index for substantia nigra (SN) and ventral tegmental area (VTA) neurons isolated from tgNM mice relative to wild-type (wt) controls at 3, 12, and 20 months of age. Pathways were identified by GSEA based on the Normalized Enrichment Score (NES). The Jaccard Index quantifies the overlap between pathway gene sets, synthesizing biological processes linked to progressive molecular changes in dopaminergic neurons.

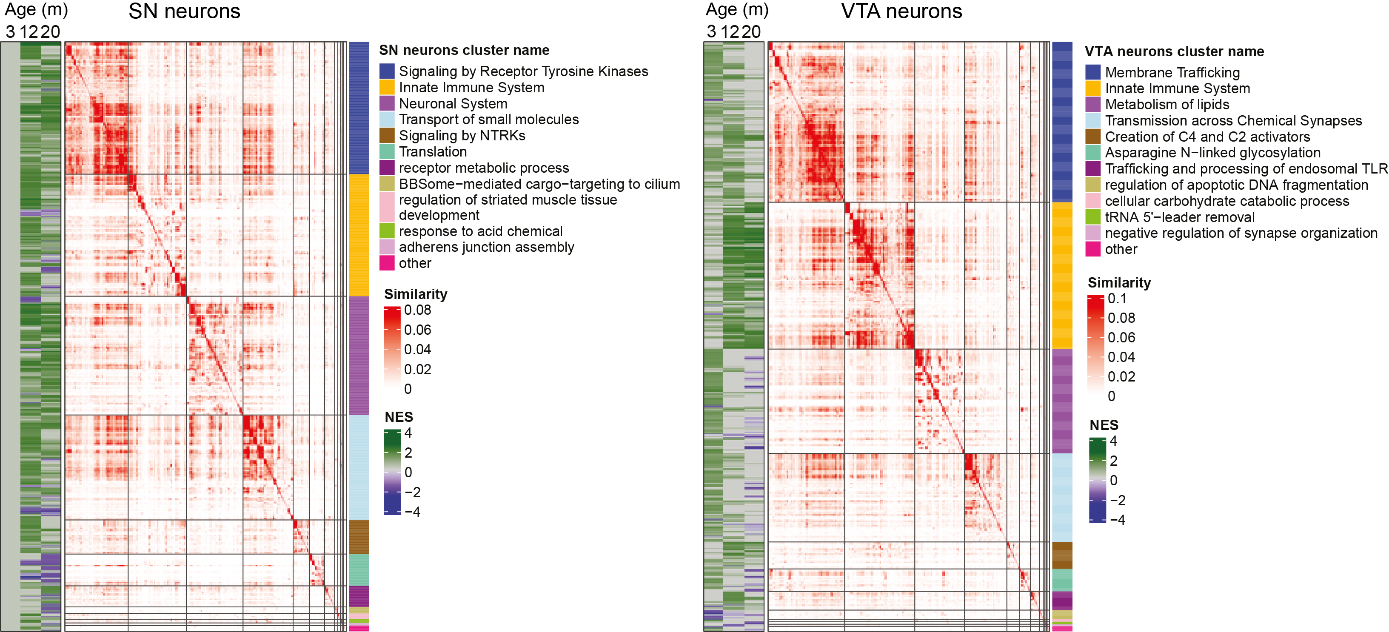

**Supplementary Figure 8. GPNMB expression in tgNM mice and PD postmortem brains.** (a) Representative image of GPNMB immunohistochemistry in the substantia nigra (SN) of wild-type (wt) mice at 20m of age showing absence of GPNMB immunoreactivity in wt mice compared to the staining observed in tgNM littermates (Figure 4b). (b) Representative combined RNAscope and immunofluorescence images from the SN of tgNM mice at 12m of age. Gpnmb transcripts (magenta) are detected in proximity to Tyrosine hydroxylase (TH)-positive (red) and NM-containing neurons identified by brightfield imaging (BF), as well as in Iba1-positive microglial cells (green). Scale bar, 10 μm. Higher magnifications from these images are shown in Figure 4c. (c) Quantification of GPNMB relative expression levels measured in human cultured melanocytes (Melan.), and different human control postmortem brain regions (substantia nigra [SN], locus coeruleus [LC], and frontal cortex [FC]). (d) Fold change (FC) in GPNMB expression in human FC from postmortem PD patients relative to control individuals. (c-d) Boxplots represent median and interquartile range with individual data points. (e) Representative images of GPNMB double-immunofluorescence staining in postmortem human SN from control (65 years, Male) and PD cases (65 years, Male). GPNMB immunoreactivity is detected in TH-positive neurons with NM (filled arrowheads) and in TH-negative neurons with NM (open arrowheads), as well as in Iba1-positive microglial cells (arrows) surrounding NM-containing cells and extracellular NM granules. Higher magnifications from these images are shown in Figure 4h. Scale bars: 20 µm

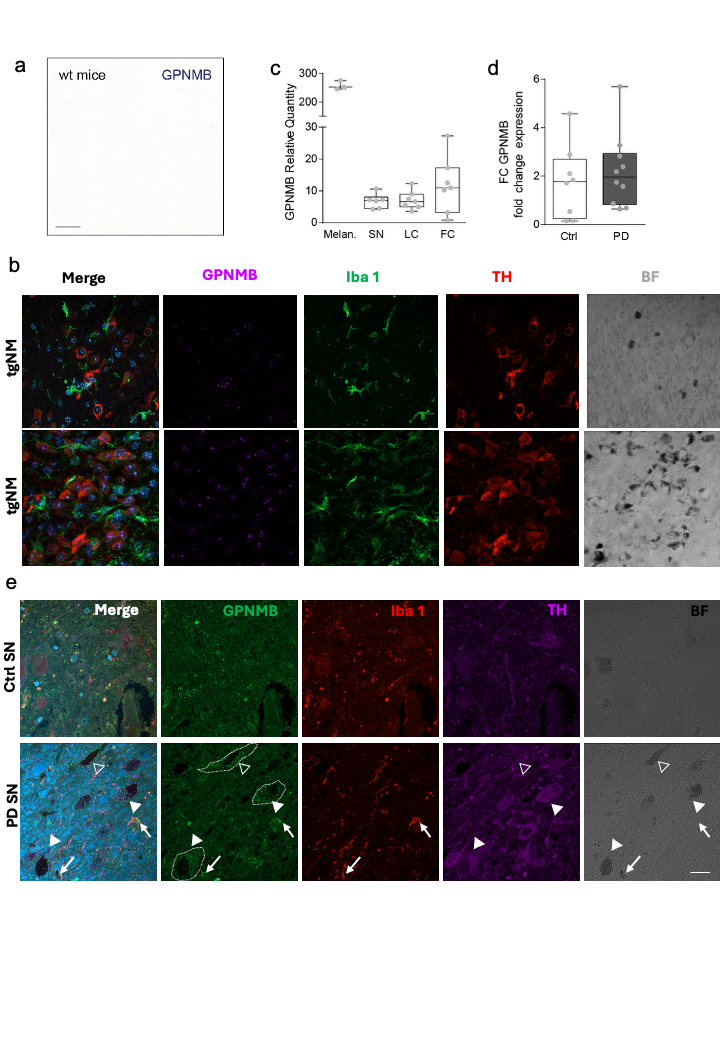

**Supplementary Table 1. PD postmortem datasets correlation with tgNM SN at different ages.** ROAST analysis comparing tgNM mice SN region (SN region, left; SN neurons, right) to the corresponding human PD transcriptomic signature from postmortem studies. Public GEO datasets used for the analysis include studies analyzing SN bulk dissections (GSE20163, GSE20164, GSE20292, GSE20333, GSE43490 and GSE7621) and LCM isolated SN neurons (GSE20141 and GSE24378) from PD patients compared to healthy controls. Mixed p-values (p) and FDR (up and downregulated DEGs) are shown. p≤0.05 are highlighted in bold and underlined.

|  | | **tgNM SN region** | | | | | | **tgNM SN neurons** | | | | | |
| --- | --- | --- | --- | --- | --- | --- | --- | --- | --- | --- | --- | --- | --- |
|  |  | 3m | | 12m | | 20m | | 3m | | 12m | | 20m | |
| PD postmortem study | Ngenes | p | FDR | p | FDR | p | FDR | p | FDR | p | FDR | p | FDR |
| GSE20141 (SN neurons) | 135 | 0.337 | 0.840 | 0.517 | 0.898 | 0.081 | 0.478 | 0.675 | 0.781 | 0.221 | 0.470 | 0.056 | 0.094 |
| GSE20163 (Bulk SN) | 86 | 0.793 | 0.907 | 0.559 | 0.898 | 0.382 | 0.478 | 0.454 | 0.749 | 0.239 | 0.470 | **0.033** | 0.093 |
| GSE20164 (Bulk SN) | 21 | 0.902 | 0.927 | 0.612 | 0.898 | 0.387 | 0.478 | 0.275 | 0.749 | 0.483 | 0.527 | 0.134 | 0.157 |
| GSE20292 (Bulk SN) | 289 | 0.503 | 0.840 | 0.511 | 0.898 | 0.172 | 0.478 | 0.447 | 0.749 | 0.271 | 0.470 | **0.044** | 0.093 |
| GSE20333 (Bulk SN) | 20 | 0.639 | 0.840 | 0.127 | 0.898 | 0.141 | 0.478 | 0.894 | 0.911 | 0.495 | 0.530 | 0.384 | 0.419 |
| GSE24378 (SN neurons) | 47 | 0.303 | 0.840 | 0.688 | 0.898 | 0.183 | 0.478 | 0.160 | 0.749 | 0.330 | 0.470 | 0.060 | 0.094 |
| GSE43490 (Bulk SN) | 375 | 0.515 | 0.840 | 0.764 | 0.898 | 0.226 | 0.478 | 0.404 | 0.749 | 0.328 | 0.470 | **0.043** | 0.093 |
| GSE7621 (Bulk SN) | 353 | 0.611 | 0.840 | 0.624 | 0.898 | 0.309 | 0.478 | 0.402 | 0.7489 | 0.211 | 0.470 | **0.029** | 0.093 |

**Supplementary Table 2.** **PD postmortem datasets correlation with tgNM LC at different ages.** ROAST analysis comparing tgNM mice LC region to the corresponding human PD transcriptomic signature from postmortem studies. Public GEO dataset from LC bulk dissection (GSE43490) from PD patients compared to healthy controls was used for the analysis. The total number of genes (Ngenes), Mixed p-values (p) and FDR (up and downregulated DEGs) are shown. p≤0.05 are highlighted in bold and underlined.

|  | | **tgNM LC region** | | | |
| --- | --- | --- | --- | --- | --- |
|  | | 3m | | 12m | |
| PD postmortem study | Ngenes | p-value | FDR | p-value | FDR |
| GSE43490 (Bulk LC) | 392 | **1.00E-04** | **1.00E-04** | **1.00E-04** | **1.00E-04** |

**Supplementary Table 3. Gene-lists linked to PD genetically and experimentally.** ROAST analysis comparing tgNM mice SN region (left) and SN neurons (right) with PD-related gene sets from different databases. The total number of genes (Ngenes), Mixed p-values (p) and FDR (up and downregulated DEGs) are shown. p≤0.05 are in bold and underlined.^[1]^

|  | | SN region | | | | | | SN neurons | | | | | |
| --- | --- | --- | --- | --- | --- | --- | --- | --- | --- | --- | --- | --- | --- |
|  |  | 3m | | 12m | | 20m | | 3m | | 12m | | 20m | |
| PD-linked dataset | N  genes^1^ | p | FDR | p | FDR | p | FDR | p | FDR | p | FDR | p | FDR |
| ClinVar Gene-Phenotype Associations | 2 | 0.476 | 0.677 | 0.770 | 0.770 | 0.392 | 0.544 | 0.801 | 0.801 | 0.349 | 0.559 | 0.076 | 0.076 |
| DISEASES Curated Gene-Disease Assocation Evidence Scores | 29 | 0.492 | 0.677 | 0.637 | 0.770 | 0.544 | 0.544 | 0.692 | 0.790 | 0.707 | 0.707 | **0.029** | 0.063 |
| DISEASES Experimental Gene-Disease Assocation Evidence Scores | 87 | 0.407 | 0.677 | 0.599 | 0.770 | 0.499 | 0.544 | 0.379 | 0.790 | 0.528 | 0.656 | 0.072 | 0.076 |
| GAD (Gene Association Database) Gene-Disease Associations | 208 | 0.531 | 0.677 | 0.445 | 0.770 | 0.178 | 0.544 | 0.382 | 0.790 | 0.264 | 0.559 | **0.028** | 0.063 |
| GWAS Catalog SNP-Phenotype Associations | 30 | 0.439 | 0.677 | 0.318 | 0.770 | 0.445 | 0.544 | 0.680 | 0.790 | 0.574 | 0.656 | **0.048** | 0.063 |
| GWASdb SNP-Disease Associations | 634 | 0.554 | 0.677 | 0.690 | 0.770 | 0.355 | 0.544 | 0.372 | 0.790 | 0.262 | 0.559 | **0.039** | 0.063 |

Correlation analyses for SN and LC with PD-associated datasets were performed at different times and therefore include different numbers of genes.

**Supplementary Table 4. Gene-lists linked to PD genetically and experimentally.** ROAST analysis comparing tgNM mice LC region with PD-related gene sets from different databases. The total number of genes (Ngenes), Mixed p-values (p) and FDR (up and downregulated DEGs) are shown. p≤0.05 are in bold and underlined.

|  | | **tgNM LC region** | | | |
| --- | --- | --- | --- | --- | --- |
|  |  | 3m | | 12m | |
| PD-linked dataset | Ngenes^1^ | p | FDR | p | FDR |
| ClinVar Gene-Phenotype Associations | 2 | **2.66E-01** | **2.66E-01** | **4.98E-01** | **4.98E-01** |
| DISEASES Curated Gene-Disease Assocation Evidence Scores | 27 | **1.00E-04** | **1.00E-04** | **4.00E-04** | **4.00E-04** |
| DISEASES Experimental Gene-Disease Assocation Evidence Scores | 92 | **1.00E-04** | **1.00E-04** | **1.00E-04** | **1.00E-04** |
| GAD (Gene Association Database) Gene-Disease Associations | 210 | **1.00E-04** | **1.00E-04** | **1.00E-04** | **1.00E-04** |
| GWASdb SNP-Disease Associations | 660 | **1.00E-04** | **1.00E-04** | **1.00E-04** | **1.00E-04** |
| GWAS Catalog SNP-Phenotype Associations | 31 | **2.00E-04** | **2.00E-04** | **3.00E-04** | **3.33E-04** |

Correlation analyses for SN and LC with PD-associated datasets were performed at different times and therefore include different numbers of genes.
